# PRDM9 directs recombination hotspots to prevent competition with meiotic transcription

**DOI:** 10.1101/2021.03.01.433380

**Authors:** Enrique J. Schwarzkopf, Omar E. Cornejo

## Abstract

PRDM9 drives recombination hotspots in some mammals, including mice and apes. Non-functional orthologs of PRDM9 are present in a wide variety of vertebrates, but why it is functionally maintained in some lineages is not clear. One possible explanation is that PRDM9 plays a role in ensuring that meiosis is successful. During meiosis, available DNA may be a limiting resource given the tight packaging of chromosomes and could lead to competition between two key processes: meiotic transcription and recombination. Here we explore this potential competition and the role that PRDM9 might play in their interaction. Leveraging existing mouse genomic data, we use resampling schemes that simulate shuffled features along the genome and models that account for the rarity of features in the genome, to test if PRDM9 influences interactions between recombination hotspots and meiotic transcription in a whole genome framework. We also explored possible DNA sequence motifs associated to clusters of hotspots not tied to transcription or PRDM9. We find evidence of competition between meiotic transcription and recombination, with PRDM9 appearing to relocate recombination to avoid said conflict. We also find that retrotransposons may be playing a role in directing hotspots in the absence of other factors.

## Introduction

Recombination, or the exchange of genetic information between homologous chromosomes, is crucial to chromosomal segregation during meiosis (Heyer et al. 2010). Since meiosis requires chromosomes to be precisely arranged while a specific set of genes are transcribed, the location of recombination events can have important consequences to the resulting gametes (Alves et al. 2017, Brick et al. 2012, da Cruz et al. 2016). Genomic regions with unusually high recombination rates (recombination hotspots) can be detrimental when inconveniently located. For instance, when mouse recombination hotspots fail to be properly directed, the resulting gametes are inviable (Brick et al. 2012). In some mammals (e.g., apes and mice), the location of recombination hotspots is associated with DNA binding sites for the zinc finger domain of the protein PRDM9 (Brick et al. 2017, Brunschwig et al. 2012, Auton et al. 2012, Myers et al. 2010, Parvanov et al. 2010). In taxa where PRDM9 is functional, it evolves quickly (Baker et al. 2017), and a multitude of hypotheses for this rapid evolution have been proposed (reviewed in Grey et al. 2018). However, the origin of PRDM9 and the reason for its functional persistence in certain taxa is still unclear. One possible explanation for the persistence of PRDM9 is that it plays a role in ensuring successful meiosis by physically separating processes competing for limited available DNA during meiosis.

During meiosis, segments of DNA must remain available for recombination and transcription to occur. How these processes interact while sharing this limited resource may affect where they act along the genome. Precise activation and inactivation of transcription is key to a highly regulated process like meiosis (da Cruz et al. 2016). At the same time, homologous recombination helps ensure the pairing of homologous chromosomes necessary for proper chromosomal segregation (Heyer et al. 2010). The occurrence of chiasmata at inappropriate locations can lead to aneuploidies, so the process of homologous recombination must be strictly regulated (Alves et al. 2017, Bugge et al. 1998, Hassold et al. 1995, Lamb et al. 1997, Robinson et al. 1998, Thomas et al. 2001). Considering the complex, tight structure assumed by chromosomes during meiosis, the potential for recombination to cause chromosomal aberrations is notable. Meiotic chromatin structure allows less DNA to be available for transcription than during the rest of the cell cycle. Reduced nucleosome availability combined with a precise regulation of the meiotic process have the potential to strongly favor the streamlining of meiotic transcription (da Cruz et al. 2016). Therefore, it is possible that meiotic transcription is competing for available DNA with recombination.

Recombination in mice is driven by the histone methylase PRDM9 (Parvanov et al. 2010). The ability of PRDM9 to trimethylate histone H3 proteins allows it to foster beneficial conditions for recombination while directing the location of recombination events (Baker et al. 2014, Brick et al. 2012, Buard et al. 2009, Diagouraga et al. 2018, Grey et al. 2011, Hayashi et al. 2005, Powers et al. 2016, Smagulova et al. 2011). In species that lack functional PRDM9, recombination hotspots tend to localize in transcription-associated regions (transcription start or stop sites and CpG islands), consistent with recombination requiring access to DNA (Auton et al. 2013, Choi et al. 2013, Hellsten et al. 2013, Singhal et al. 2015, Schwarzkopf et al. 2020). Additionally, recombination hotspots in PRDM9 knock-out mice form at histone methylation marks similar to those made by PRDM9 but associated with promoters and enhancers. However, unlike in species with non-functional copies of PRDM9, relocation of recombination hotspots leads to infertility in mice due to improper chromosomal segregation. This led to the suggestion that PRDM9 may decouple transcription and recombination (Brick et al. 2012).

Transposable elements (TEs) are negatively correlated with overall recombination rates, but in mice and humans, certain families of TEs are common in recombination hotspot rich regions (Kent et al. 2017, Myers et al. 2008, Campos-Sánchez et al. 2016). Some of these TEs, like L2 elements and THE1A/B retrotransposons, that are enriched near recombination hotspots contain PRDM9-binding DNA sequence motifs, which can explain the correlation (Myers et al. 2008). However, in other cases [e.g., endogenous retroviruses (ERVs) in mice], the explanation is less clear, although a high rate of recombination playing a role in mouse ERV integration has been proposed (Campos-Sánchez et al. 2016). While the relationship between specific TE families and recombination hotspots is likely overshadowed by that of PRDM9, it may be possible to find patterns when looking at certain subsets of mouse recombination hotspots.

In this study, we explored the physical relationship between recombination hotspots, transcription during meiosis, and PRDM9 in mice (*Mus musculus*). We integrated existing data sets detailing recombination hotspots (Brunschwig et al. 2012), nucleosome availability (Maezawa et al. 2018), gene expression during meiosis (Da Cruz et al. 2016), and a map of PRDM9 binding sites (Walker et al. 2015) to answer the following questions: (i) Is there competition between recombination and meiotic transcription in mice? (ii) If so, what role does PRDM9 play in this interaction? (iii) Are there DNA sequence motifs associated with recombination hotspot clusters not explained by transcription or PRDM9?

## Materials and Methods

### Determining regions of interest for analyses

In order to establish regions that were undergoing transcription during meiosis, we used data from Da Cruz et al. 2016. Specifically, we obtained transcriptomic data for mouse pachytene spermatocytes. We retained genes with least 2 Reads Per Kilobase of transcript, per Million mapped reads (RPKM), in concordance with the methods of Da Cruz et al. (2016). The locations of genes that met the cutoff were then collected in a BED file for later analyses.

We obtained recombination hotspots from available data from an LD-based recombination map for *Mus musculus* (Brunschwig et al. 2012). The locations for the hotspots were converted from build 37 (NCBI37/mm9) to build 38 (NCBI38/mm10) of the mouse genome using the *liftOver* online tool from UCSC Genome Browser (Kent et al. 2002). The minimum ratio of bases that must map was set to 0.75 and 38 hotspots were lost during the conversion. The remaining hotspot locations were then converted to the bed file. Because these data did not include hotspots on the sex chromosomes, we limited our analysis of all features to the 19 mouse autosomes.

ATAC-seq data from Maezawa et al. 2018 was downloaded from the Gene Expression Omnibus series GSE102954. Specifically, samples GSM2751129 and GSM2751130 were downloaded. These samples correspond to pachytene spermatocytes from wild-type mice.

Afinity-Seq data from Walker et al. 2015 was used to define DNA regions with affinity to the PRDM9 protein. The regions were converted from build 37 (NCBI37/mm9) to build 38 (NCBI38/mm10) of the mouse genome using the *liftOver* online tool from UCSC Genome Browser (Kent et al. 2002). The minimum ratio of bases that must map was set to 0.75 and 1 site was lost during the conversion.

### Comparing feature locations along the genome

We compared the overlaps between the various features by using a genome resampling routine. The resampling routine uses the *shuffle* command in BEDtools (version 2.27.1, Quinlan and Hall 2010) to rearrange features of the same size and number as the real ones along the mouse genome. For each pair of features, we shuffled one of the features (hotspots, available DNA, meiotic transcription, and PRDM9 binding sites) ten thousand times to generate a distribution of expected overlap sizes. We then compared the generated distribution to the observed overlap size. In this way we determined the probability of the observed overlap size under null conditions. The count of each feature is reported in Table 1. We also complemented our available DNA data to obtain a bed file of DNA bound to nucleosomes, which we compared with hotspots in order to determine if the number of hotspots that overlap with nucleosomes is more or less than expected by chance.

**Table 1.** Count and size of features used in location comparisons.

|  | Genes | Transcribed genes | Hotspots | Nucleosome availability | PRDM9 affinity |
| --- | --- | --- | --- | --- | --- |
| Count | 54547 | 7236 | 47030 | 22943 | 31838 |
| Mean (median) feature size in bp | 24814.45 (3805) | 49177.26 (25012.5) | 8235.286 (5000) | 852 (1488.5) | 320.99 (297) |
| Min-max feature size in bp | 9-4434881 | 353-2244856 | 1938-971000 | 1-10668 | 187-5409 |

### Modeling Recombination hotspots

In order to better describe how PRDM9 and meiotic transcription interact with each other and with hotspot location, we modeled hotspot counts using a Zero Inflated Poisson (ZIP) regression. We used ZIP regressions because of the high proportion of the genome that did not include any of our features of interest, leading to an excess of zeros in our data. We applied the ZIP regression using the *zeroinfl* function from the *pscl* (version 1.5.5; Zeileis et al. 2008) package in *R* (version 3.6.1; R Core Team 2017) on the *Mus musculus* genome by breaking it up into 20 kilobase (kb) windows. This window size was selected to allow for the possibility of multiple hotspots in each window and therefore model a hotspot count. In each window we counted the number of PRDM9 affinity sites, meiotically transcribed genes, and recombination hotspots. These features were still often absent in the windows, leading to an excess of zeros. We initially used a Generalized Linear Model (GLM), run with the *glm* function from the *stats* package in R (version 3.6.1; R Core Team 2017). The excess zeros made the GLM unsuccessful and ZIP regressions the appropriate choice.

To further test the role of PRDM9 and meiotic transcription in the location of hotspots along the genome we used our knowledge from the ZIP regression to fit a Zero Inflated Poisson Hidden Markov Model (ZIPHMM). Using a ZIPHMM we are able to describe the hotspot counts in windows with respect to the hotspot counts in the preceding window and to the number of meiotically transcribed genes and PRDM9 affinity regions. We obtained counts from 20 kb non-overlapping windows along the *Mus musculus* genome and used the *hmmfit* function from the *ziphsmm* package (version 2.0.6; Xu and Liu 2018) in *R* (version 3.6.1; R Core Team 2017) to fit a ZIPHMM model with PRDM9 affinity region count as a covariate for structural zeros and meiotically transcribed genes count as a covariate for the state. With the parameters for the ZIPHMM established we estimated the likely state of each window and describe the states in the ZIPHMM. With the state descriptions and their distribution along the genome we then drew conclusions about the relationship between PRDM9 affinity regions, meiotically transcribed genes, and recombination hotspots. The structure of the ZIPHMM is described in Supplementary Figure 1.

### Exploring DNA sequence motifs in unexplained hotspot clusters

In order to establish if there are DNA sequence motifs associated with recombination hotspot clusters not associated with transcription or PRDM9 affinity, we used RepeatMasker (Smit et al., 2013) to search for DNA sequence motifs in windows that are rich in recombination hotspots but poor in PRDM9 affinity sites and meiotically transcribed genes. We chose windows with at least four hotspots, zero PRDM9 affinity sites, and zero meiotically transcribed genes. Additionally, we removed windows with PRDM9 affinity sites or transcribed genes in immediately neighboring windows. We ran RepeatMasker using the slow option (−s) and we chose mouse as the species. We also ran RepeatMasker on the entire mouse genome fasta to establish general genome expectations to compare to our findings in specific windows.

## Results

### Comparing feature locations along the genome

Recombination hotspots are overrepresented in nucleosome depleted regions (Figure 1A). They, therefore, appear to require access to DNA in the form of nucleosome depletion. However, observed recombination hotspots that overlapped with a nucleosome were 14.67 times the average number from simulations, indicating an overrepresentation of nucleosomes in recombination hotspots. This is consistent with findings that recombination hotspots in mice often have a nucleosome embedded in them but are otherwise in nucleosome-free regions (Getun et al. 2010, Getun et al. 2012). Conversely, we did not find evidence of a clear overrepresentation or underrepresentation of recombination hotspots near transcribed genes (Figure 1B, Supplementary Table 1). Yet, when we subset hotspots by their proximity to a region known to have affinity to PRDM9, PRDM9 hotspots were underrepresented near meiotically transcribed genes (Figure 1C-D, Supplementary Table 1). This trend appears unique to meiotically transcribed genes, as identical analyses using all annotated mouse genes found hotspots to be overrepresented regardless of PRDM9 affinity (Supplementary Table 1).

**Figure 1.**
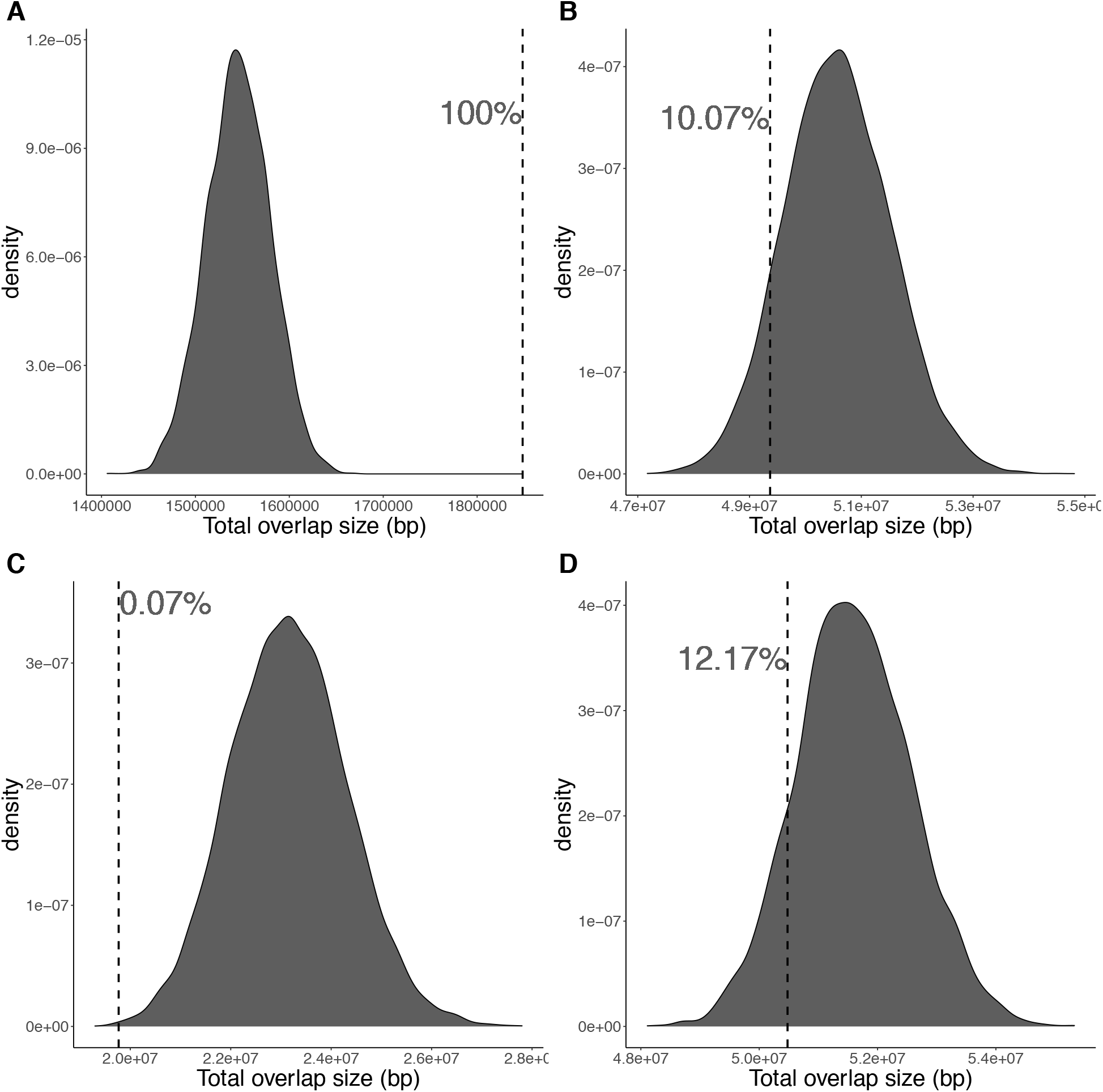
Expected distribution of overlap sizes based on shuffling features along the mouse genome. Dashed vertical lines represent the location of observed overlap sizes in the distributions of simulated overlap sizes and they are labeled with the percentage of expected overlap sizes that are less than the observed. The features compared are: (A) recombination hotspots and nucleosome availability, (B) recombination hotspots and meiotically transcribed genes, (C) recombination hotspots associated with a PRDM9 affinity region and meiotically transcribed genes, and (D) recombination hotspots not associated with a PRDM9 affinity region and meiotically transcribed genes.

### Modeling recombination hotspots

To further explore the relationship between meiotic transcription, PRDM9 affinity, and recombination hotspots, we modelled their counts in 20kb windows along the genome using a Zero Inflated Poisson (ZIP) regression. Our best fit ZIP regression was one where increasing the number of PRDM9 affinity regions lowers the probability that the window has zero hotspots and increasing in the number of meiotically transcribed genes reduces the number of hotspots in the window (Table 2). For comparison and internal validation of the approach, we also modeled the number of hotspots in a window using a Generalized Linear Model (GLM) but found that the best fit GLM was not as good of a fit for our data than the ZIP regression (Supplementary Table 2).

**Table 2.** Parameters of the best fit zero-inflated Poisson (ZIP) regression model of the hotspot count in 20kb windows across all 19 mouse autosomes. The Transcription variable is the number of transcribed genes in a window and the PRDM9 variable is the number of PRDM9 affinity regions in the window. This model had an AIC=242442.3.

| Model portion | Parameter | Estimate | Std. Error | p-value |
| --- | --- | --- | --- | --- |
| *Count model* | Intercept | -0.225760 | 0.006884 | <2e-16 |
|  | Transcription | -0.037574 | 0.009610 | 9.23e-05 |
| *Zero-inflation model* | Intercept | -0.64619 | 0.01772 | <2e-16 |
|  | PRDM9 | -0.45925 | 0.02986 | <2e-16 |

Since the ZIP regression considers each window as independent from the rest, we also modeled the windows with a ZIP Hidden Markov Model (ZIPHMM) which accounts for neighboring windows when predicting the number of hotspots in the focal window. We constructed ZIPHMM models where for each window, the covariate for structural zeros was the number of PRDM9 affinity regions and the covariate for the number of hotspots was the number of meiotically transcribed genes. Our best fit ZIPHMM model included two states, which generally corresponded with absence (State 1) and presence (State 2) of hotspots (Table 3). The ZIPHMM had an AIC lower than that of the best fit ZIP regression (ZIPHMM AIC = 175165.65, ZIP AIC = 242442.3), indicating that the state of the previous window is important to the prediction of hotspot presence in a window (Figure 2, Table 4, Supplementary Figures 2–19).

**Table 3.** Feature counts in windows identified as each of the two states of the best fit ZIPHMM. The number of windows in a state with a particular feature count are presented in each cell along with the percentage of windows from that state that have that particular feature count (in parentheses).

|  |  | **Count** |  |  |  |  |  |
| --- | --- | --- | --- | --- | --- | --- | --- |
| **Feature** | **State** | **0** | **1** | **2** | **3** | **4** | **5** |
| *Hotspots* | 1 | 64684<br>(91.3%) | 5871<br>(8.3%) | 321<br>(0.5%) | 6<br>(0.008%) | 0<br>(0%) | 0<br>(0%) |
|  | 2 | 12695<br>(24.3%) | 23539<br>(45%) | 12373<br>(23.7%) | 3201<br>(6.1%) | 439<br>(0.8%) | 19<br>(0.04%) |
| *PRDM9* | 1 | 56074<br>(79.1%) | 12909<br>(18.2%) | 1747<br>(2.5%) | 139<br>(0.2%) | 13<br>(0.02%) | 0<br>(0%) |
|  | 2 | 39813<br>(76.2%) | 10826<br>(20.7%) | 1468<br>(2.8%) | 149<br>(0.2%) | 8<br>(0.02%) | 2<br>(0.004%) |
| *Transcription* | 1 | 57569<br>(81.2%) | 12119<br>(17%) | 1086<br>(1.5%) | 90<br>(0.1%) | 18<br>(0.03%) | 0<br>(0%) |
|  | 2 | 42895<br>(82.1%) | 8458<br>(16.2%) | 808<br>(1.5%) | 90<br>(0.2%) | 14<br>(0.03%) | 1<br>(0.002%) |

**Figure 2.**
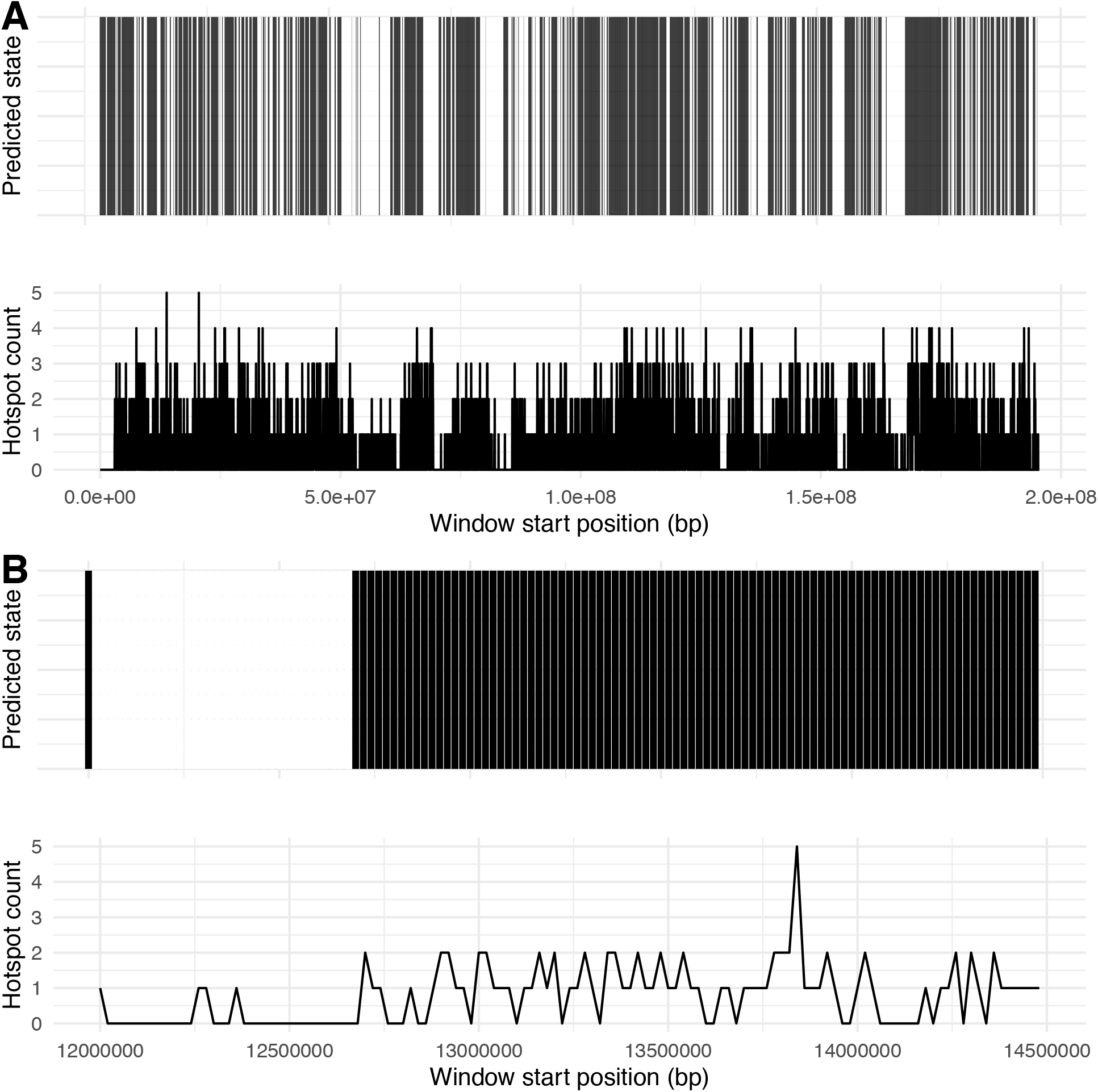
Predicted states according to best fit zero-inflated Poisson hidden Markov model (ZIPHMM) and hotspot count for all chromosome 1 windows (A) and for a small section of chromosome 1 (B).

**Table 4.** Summary of the ZIPHMM’s ability to predict the presence of hotspots (HS in the table). A true negative is defined as a window with no hotspots being assigned to state 1 and a true positive is defined as a window with at least one hotspot being assigned to state 2.

|  | State = 1 | State = 2 | Total |  |
| --- | --- | --- | --- | --- |
| HS count = 0 | 64684 | 12695 | 77379 | *True negative rate = 83.59%* |
| HS count > 0 | 6198 | 39571 | 45769 | *True positive rate = 86.46%* |
| Total | 70882 | 52266 | 123148 |  |
|  | *Specificity = 91.26%* | *Sensitivity = 75.71%* |  |  |

### Exploring DNA sequence motifs in unexplained hotspot clusters

We explored possible DNA sequence motifs in hotspot rich windows in genomic regions lacking PRDM9 affinity sites and meiotically transcribed genes. These windows had an abundance of mammalian long terminal repeat retrotransposons (ERVL-MaLRs; Table 5). In mice, ERVL-MaLRs likely still expand, indicating the persistence of a reverse transcriptase (Smit 1996). We also observed a lack of Alu/B1 elements in these windows (Table 5). Notably, Alu/B1 are unusual in that they are SINE elements without a polymerase III promoter (Smit 1996).

**Table 5.**
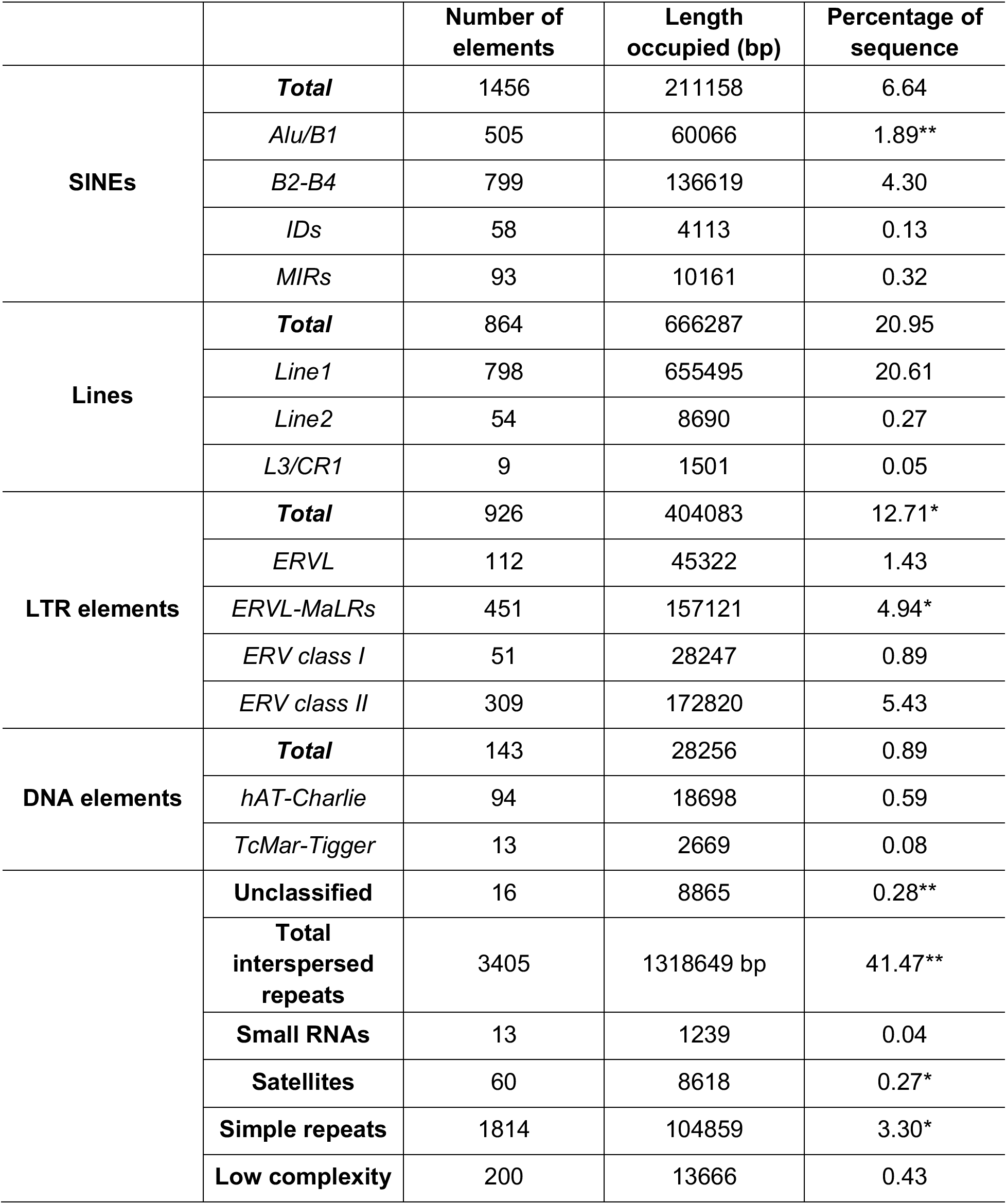
Summary of RepeatMasker results for windows with at least 4 hotspots, no PRDM9 affinity regions, and no meiotically transcribed genes. Selected windows also had no PRDM9 affinity regions or meiotically transcribed genes in their flanking windows. Values greater than the 95% quantile of resampled distribution are denoted by *. Values less than the 5% quantile of resampled distribution are denoted by **.

## Discussion

### Recombination hotspots along the mouse genome are in nucleosome depleted regions

In order to identify competition for available DNA between recombination and meiotic transcription, it was important that we first confirm our assumption that recombination requires available DNA. We found that recombination hotspots are located in regions mostly, but not completely depleted of nucleosomes. This is consistent with hotspots usually being in nucleosome-free regions that are either flanked by nucleosomes or occasionally have an embedded nucleosome. This pattern of nucleosome distribution has been observed for specific mouse recombination hotspots (Getun et al. 2010, Getun et al. 2012). We found evidence for this pattern being consistent throughout the mouse genome. Confirming the genome-wide consistency of this pattern by analyzing each hotspot individually was inappropriate given the varied origin of our data. Because the data does not reflect a single population, specific relationships are more likely to be due to random chance. Nevertheless, the detection of these broad-scale patterns consistent with previous findings provides evidence consistent with our assumption for the need of available DNA for recombination.

### PRDM9 associated hotspots are decoupled from meiotic transcription

Multiple lines of evidence supported our hypothesis that PRDM9 decouples meiotic transcription from recombination hotspots. First, recombination hotspots that overlap with PRDM9 affinity sites are unlikely to be near meiotically transcribed genes. Second, our best-fit ZIP regression model indicates that the number of PRDM9-associated hotspots in a given genomic region is inversely proportional to the number of meiotically transcribed genes. To further establish the consistency of these results we looked at a set of genes that have been identified to be essential to meiosis. We found that the kinetochore protein coding gene, *Ndc80*, which is a molecular switch controlling kinetochores assembly, is expressed during pachytene spermatogenesis (Chen et al. 2017), does not overlap with a recombination hotspot. This is what we expected for a gene that can severely impact meiosis, and it’s what we observed for another such gene, the spermatogenesis transcription factor *Sox30* (Chen et al. 2018). *Sox30*-knockout mice show evidence of arrested early spermatid development (Chen et al. 2018). Independent studies have confirmed that *Sox30* is expressed almost uniquely during formation of meiotic spermatocytes and postmeiotic haploids (Bai et al. 2018), illustrating the importance of timing in the displacement of recombination and essential transcription. Crucially, *Sox30* initiates expression during early meiotic cell stages but controls a core program of gene expression that extends beyond post-meiosis (Bai et al. 2018). *Prm1, Tnp1*, and *Tnp2*, which all play a role in chromatin shaping during meiosis (da Cruz et al. 2016), are also expressed during meiosis but do not overlap with recombination hotspots. These results are particularly important because abnormal or extemporal expression of *Tnp* and *Prm* genes have been shown to cause infertility and spermiogenesis arrest (Lee et al. 1995, Tseden et al. 2007, Francis et al. 2014). A common feature of the genes discussed above is that the timing of their expression is crucial to the normal programming of meiosis. Future work should focus on disentangling which essential genes are transcriptionally restricted to a particular window of time, during the pachytene stage.

Recently, the loss of *Fbxo47* in mammalian cells has been found to result in a breakdown of meiotically important, telomere-associated processes and tied to infertility in male mice (Chen et al. 2018, Hua et al. 2019). Yet, we find that, though it is expressed during meiosis I in mice, its location overlaps with a recombination hotspot. However, we found that the recombination hotspot associated with this *Fbxo47* is located at the end of the coding sequence, which may reduce any conflict between recombination and transcription. We also observed that some meiotically expressed members of the *Fos* family of genes do overlap with hotspots of recombination. This finding is consistent PRDM9 contributing to minimize the possibility for physical conflicts between transcription of essential genes during meiosis and successful recombination, due to the possibility of a “rescue effect” by other genes of the family.

Collectively, these observations further support that the decoupling of recombination and transcription of essential meiotic genes is necessary for successful meiosis to occur, emphasizing the importance of PRMD9. PRDM9 is also a known hybrid sterility gene in mice, producing infertility in hybrids of mouse strains with different PRDM9 alleles (Mihola et al. 2009). This sterility is driven by asynapsis between chromosomes from the two differing strains and can be reversed when long tracts of between-strain homology are inserted into the more problematic chromosomes (Gregorova et al. 2018). Additionally, human PRDM9 often binds to promoter regions and can activate the expression of some genes (Altemose et al. 2017). Such a role has not been identified in mouse, though methylation marks similar to those produced by PRDM9 are found in mouse promoter regions during meiosis (Brick et al. 2012). When PRDM9 is knocked-out in mice, however, these methylation marks remain while those outside of promoter regions are lost (Brick et al. 2012). Furthermore, double-strand break hotspots are found at promoter regions, indicating that PRDM9 redirects recombination away from functional genomic elements (Brick et al. 2012). Our findings further indicate that PRDM9 specifically redirects recombination away from meiotically transcribed genes. Given that PRDM9 knock-out mice are completely sterile, PRDM9 may be maintained due to a relaxation of other controls on recombination at meiotically important promoter regions. This would lead to a loss of the ability to properly segregate chromosomes in the absence of a protein that drives recombination away from regions involved in meiotic transcription.

The recombination-directing function of PRDM9 has been lost multiple times across vertebrates (Baker et al. 2017). This loss of function has not precluded the formation of recombination hotspots, rather they are relocated to transcription-associated regions (Auton et al. 2013, Singhal et al. 2015). Why some species can tolerate the loss of PRDM9 when it produces infertility in mice is unclear (Brick et al. 2012). One possible explanation, given our finding that PRDM9 decouples meiotic transcription from recombination and that methylation markers similar to those produced by PRDM9 are found at promoter regions in PRDM9 knock-out mice (Brick et al. 2012), is that mice have a more streamlined meiotic epigenome. A streamlined epigenome may limit the number of available targets for recombination in the absence of PRDM9, leading to the transcription-recombination conflict. Recent work on snake recombination found support for PRDM9 as a driver of recombination hotspots (Schield et al. 2020). However, snake recombination hotspots are still found near functional genomic elements, as is the case in non-PRMD9 species. One possible explanation is that snake PRDM9 is methylating promoter regions of genes not transcribed during meiosis. These methylation marks, like those in mice, likely do not initiate transcription (Brick et al. 2012). Rather, they provide additional targets for recombination, preventing the conflict between recombination and transcription. Human PRDM9 is known to bind to promoter regions and in some cases activate transcription (Altemose et al. 2017), which points to a potential origin of PRDM9 as a transcription factor regulator that was later co-opted to regulate recombination. In species that lack PRMD9, hotspots of recombination seem to be associated to promoter regions and would also explain who PRDM9 could have evolved into a regulator of recombination. It is possible that snake PRDM9 plays a similar role in binding to promoter regions but methylates them in a way that favors recombination rather than transcription. The dynamics of snake PRDM9 may serve as an precursor to the loss of PRDM9, potentially giving way to a more relaxed meiotic epigenome at which point the loss of PRDM9 no longer imposes such a steep cost to fitness. If a streamlined epigenome is the main difference between PRDM9 knock-out tolerant and intolerant species, we expect that the conflict between recombination and meiotic transcription would be driven by transcription preventing proper recombination rather than the other way around.

### Unexplained recombination hotspot clusters tied to retrotransposons

If PRDM9 is acting to decouple recombination and transcription, but not all recombination hotspots are associated with a PRDM9 affinity site, then what could explain the location of these unexplained hotspots? Regions with no meiotically transcribed genes that contain clusters of non-PRMD9 hotspots are also associated with TEs that encode a reverse transcriptase (Table 5, Smit 1996). This indicates that non-PRDM9 hotspots in mice are associated with some form of transcription, whether that be meiotic transcription or TE activity. Mouse ERVs in general have been observed to be overrepresented in hotspot rich regions (Cuevas-Sánchez et al. 2016). ERVL-MaLRs specifically, however, may act as an auxiliary mechanism for the separation between recombination and meiotic transcription. Expression of the reverse transcriptase encoded in ERVL-MaLRs likely generates favorable conditions for recombination to occur, without incurring an evolutionary cost to the host if it conflicts with recombination. These retrotransposon-rich regions are also most likely evolutionarily neutral and therefore insensitive to frequent retrotransposon activity (Batzer and Cordaux 2009). That activity, specifically when it happens during meiosis, could promote higher rates of recombination. While ours is a simple approach and only provides evidence of a correlations between TE activity and recombination hotspots in mice, it does invite further study in a variety of species to elucidate the history of this relationship.

### Potential caveats

Combining existing datasets to answer questions that they were not originally intended to answer comes with a number of challenges. One that we were keenly aware of is the differences in evolutionary history between the organisms used in the different studies. While we were careful to ensure that all the studies we used were from the same or similar lab mouse strains, it’s inevitable that there will be differences in their genetic history. These differences can bias our results, though we expect that they will generally obscure existing relationships. Therefore, we expect that the relationships we are able to detect are likely true, if incomplete.

### Concluding remarks

We explored the potential for interactions among factors that influence the location of recombination hotspots. We found additional support for the previously studied hypotheses that PRDM9 relocates recombination to avoid transcription-associated regions. We further proposed and found evidence for the decoupling of recombination and transcription by PRDM9 being specific to meiotic transcription. As genomic resources expand both in quantity and in variety, leveraging their information can facilitate exploration of new hypotheses. Using existing data is not without drawbacks, but we expect that the relationships we identified are strong enough to overcome those drawbacks. Leveraging existing genomic data is an efficient way to explore complex questions about broad patterns in the genome.

---

**Supplementary table 1.** Percentage of simulated overlaps that were smaller than the observed overlap for pairs of features.

|  | **Meiotically transcribed genes** | **All genes** |
| --- | --- | --- |
| **Hotspots** | 10.07 % | 100 % |
| **Hotspots-PRDM9** | 12.25 % | 87.79 % |
| **Hotspots+PRDM9** | 0.07 % | 100 % |

**Supplementary table 2.**
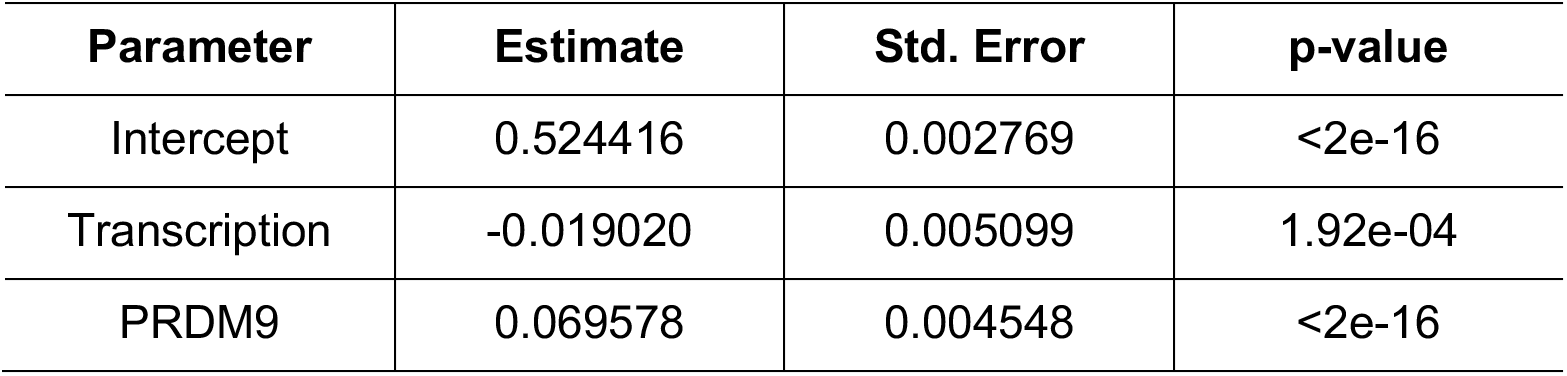
Parameters of the best fit GLM to predict the number of hotspots in 20kb windows across all 19 mouse autosomes. The Transcription variable is the number of transcribed genes in a window and the PRDM9 variable is the number of PRDM9 affinity regions in the window. This model had an AIC=297477.

| **Parameter** | **Estimate** | **Std. Error** | **p-value** |
| --- | --- | --- | --- |
| Intercept | 0.524416 | 0.002769 | <2e-16 |
| Transcription | -0.019020 | 0.005099 | 1.92e-04 |
| PRDM9 | 0.069578 | 0.004548 | <2e-16 |

**Supplementary figure 1.**
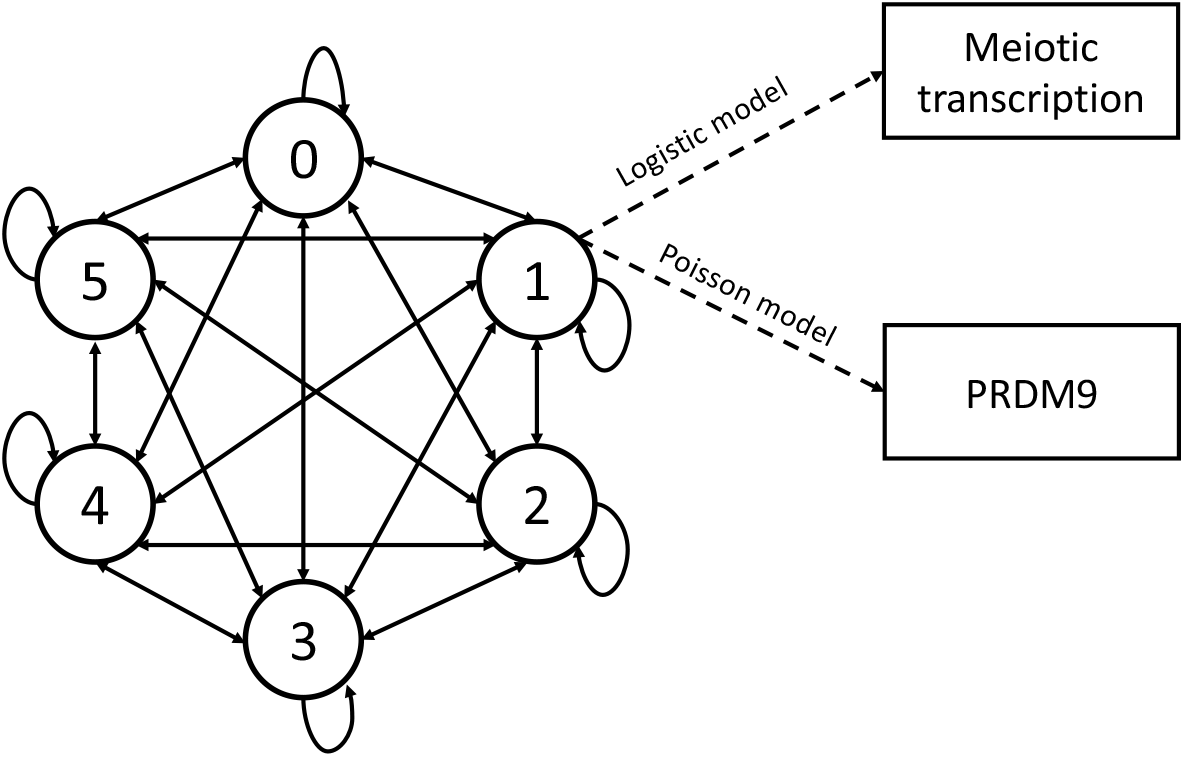
A representation of our zero-inflated Poisson hidden Markov model (ZIPHMM). Circles represent the different states of the Markov chain. The number in the circle is the number of recombination hotspots in each window. The dotted lines represent the emission probabilities. There are different emission probabilities for each state in the Markov chain. For simplicity, only one is represented here. The rectangles represent the observed sets of states. Each rectangle represents a set of possible states (e.g. PRDM9 can take count values between 0 and 5).

**Supplementary figure 2.**
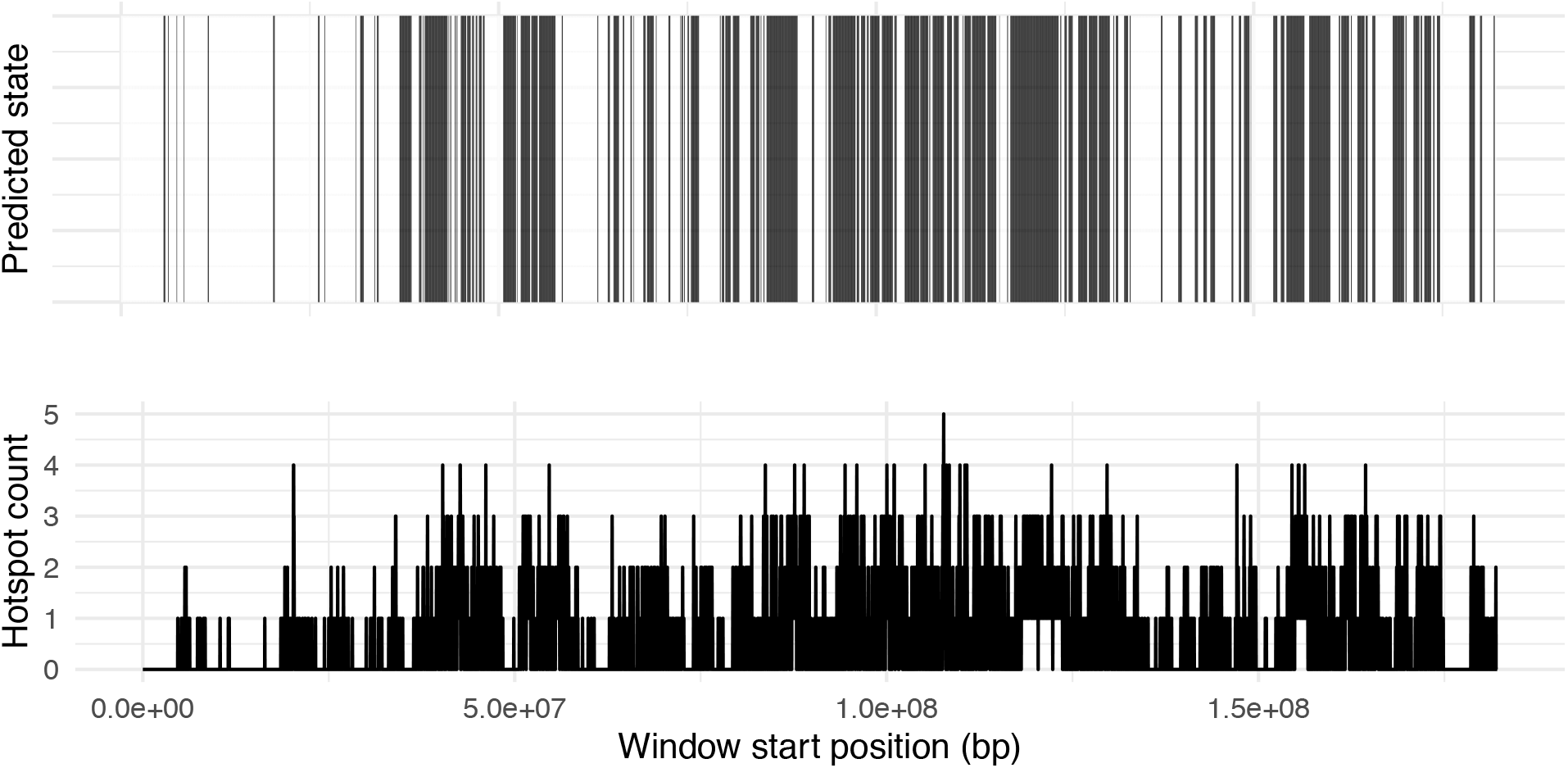
Predicted states according to best fit ZIPHMM and hotspot count for all chromosome 2 windows.

**Supplementary figure 3.**
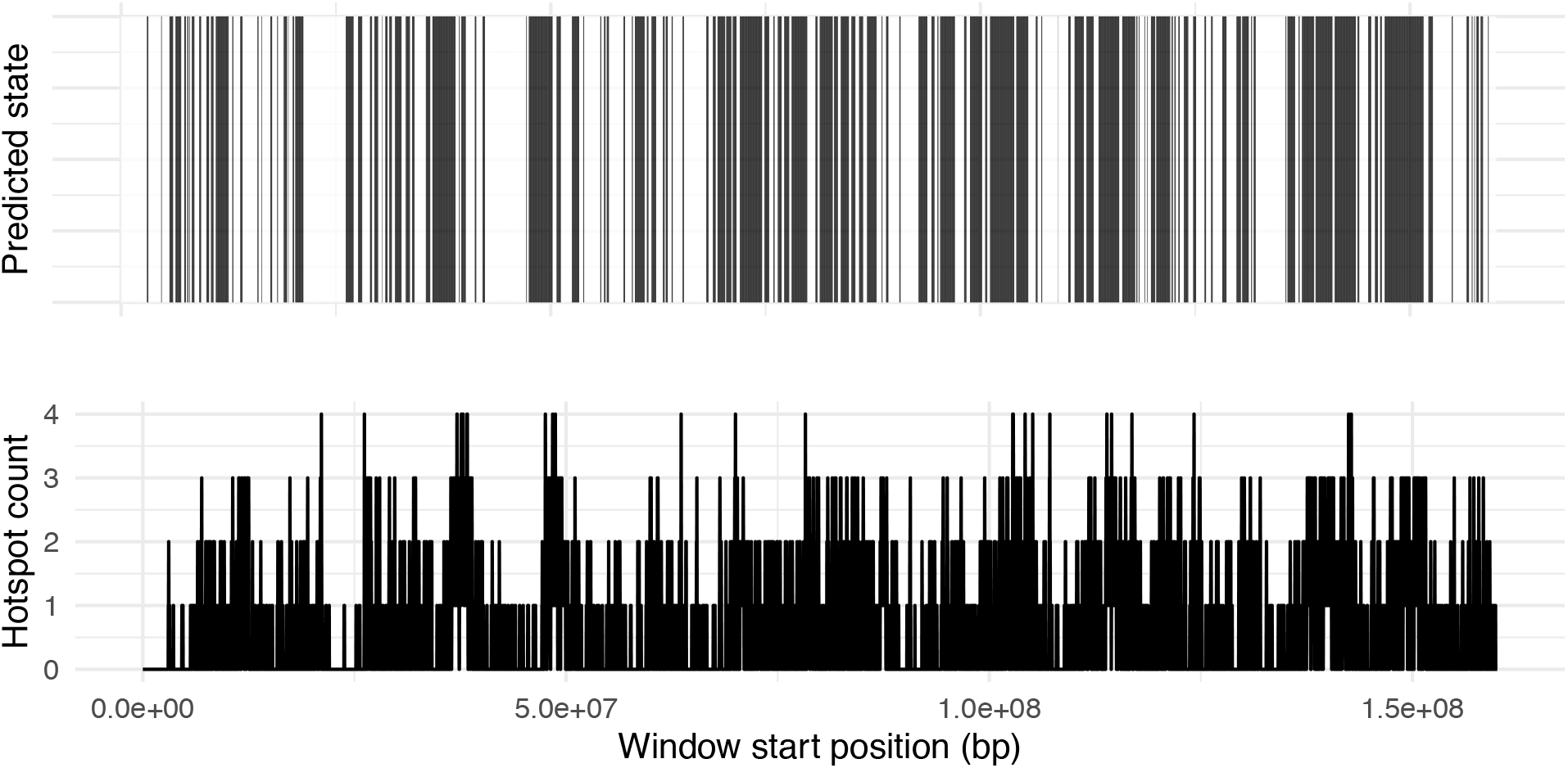
Predicted states according to best fit ZIPHMM and hotspot count for all chromosome 3 windows.

**Supplementary figure 4.**
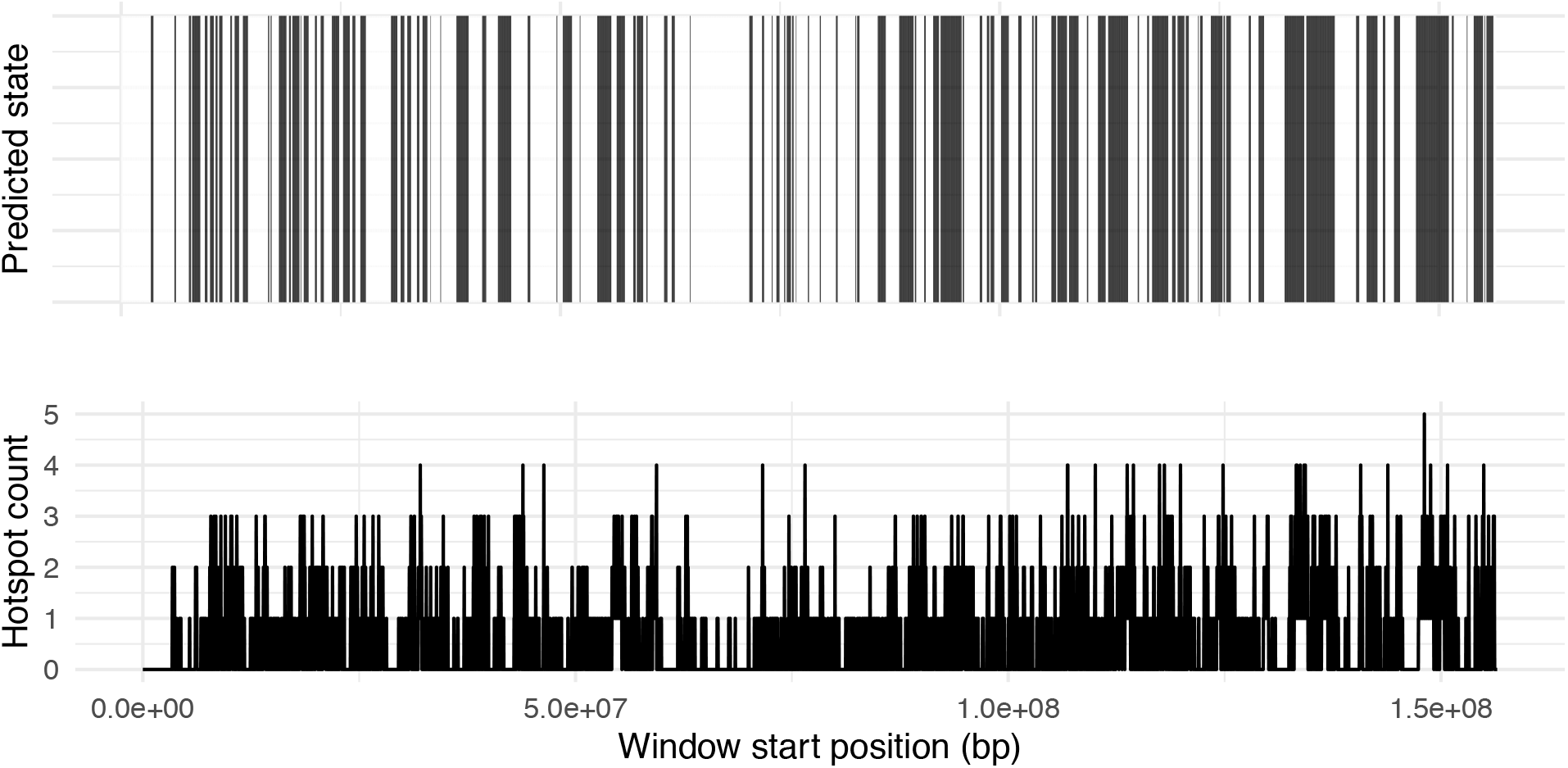
Predicted states according to best fit ZIPHMM and hotspot count for all chromosome 4 windows.

**Supplementary figure 5.**
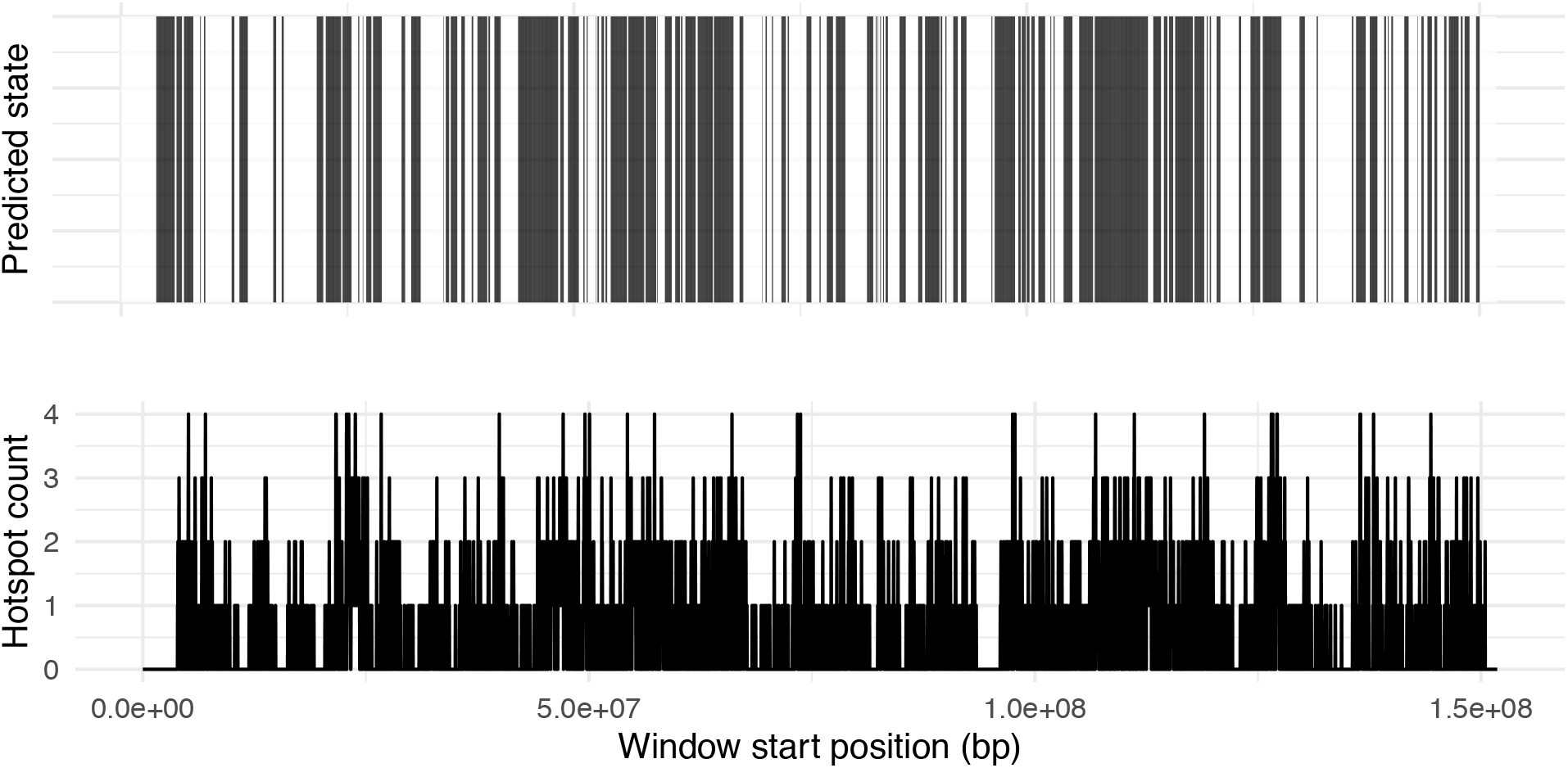
Predicted states according to best fit ZIPHMM and hotspot count for all chromosome 5 windows.

**Supplementary figure 6.**
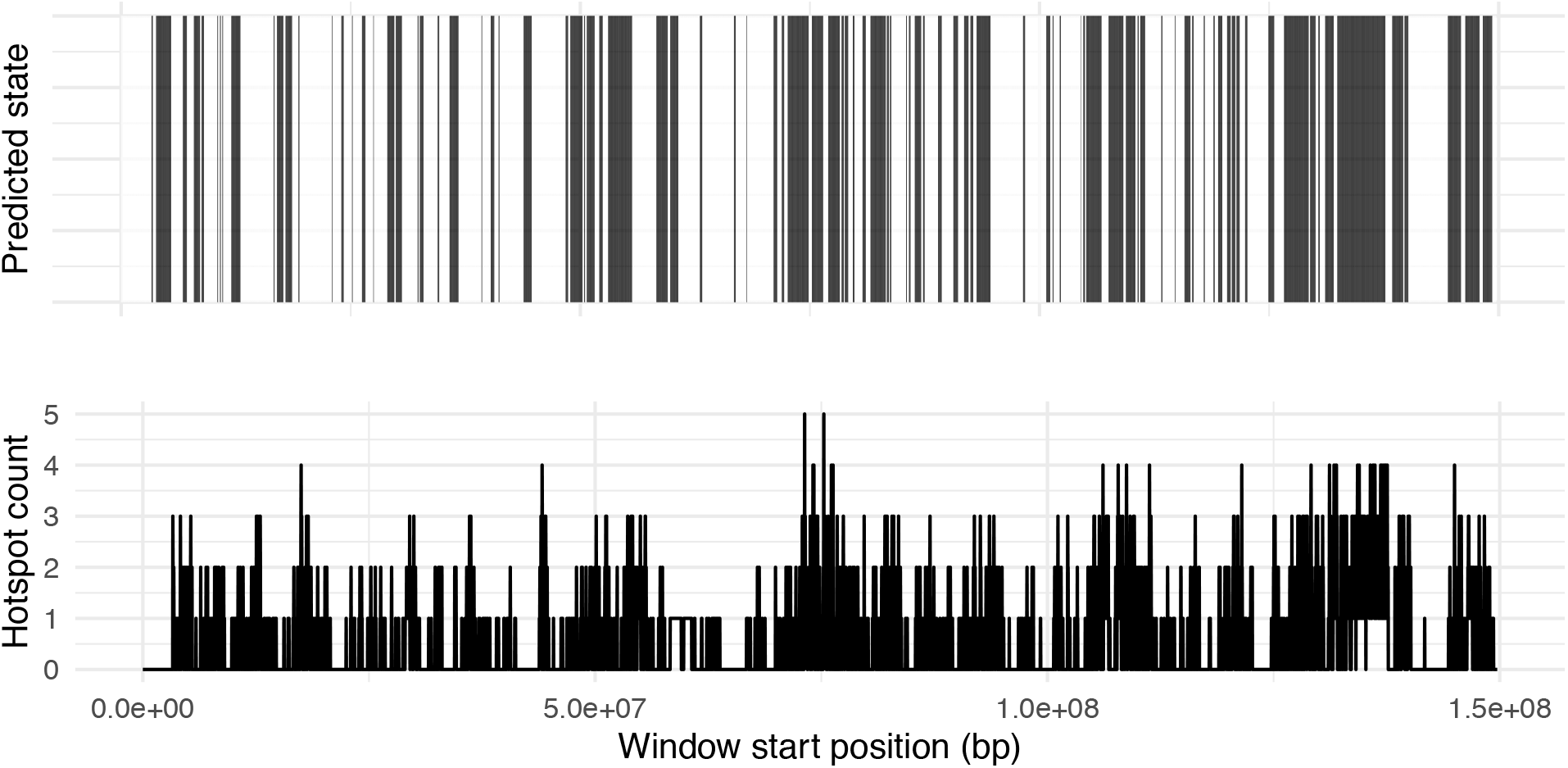
Predicted states according to best fit ZIPHMM and hotspot count for all chromosome 6 windows.

**Supplementary figure 7.**
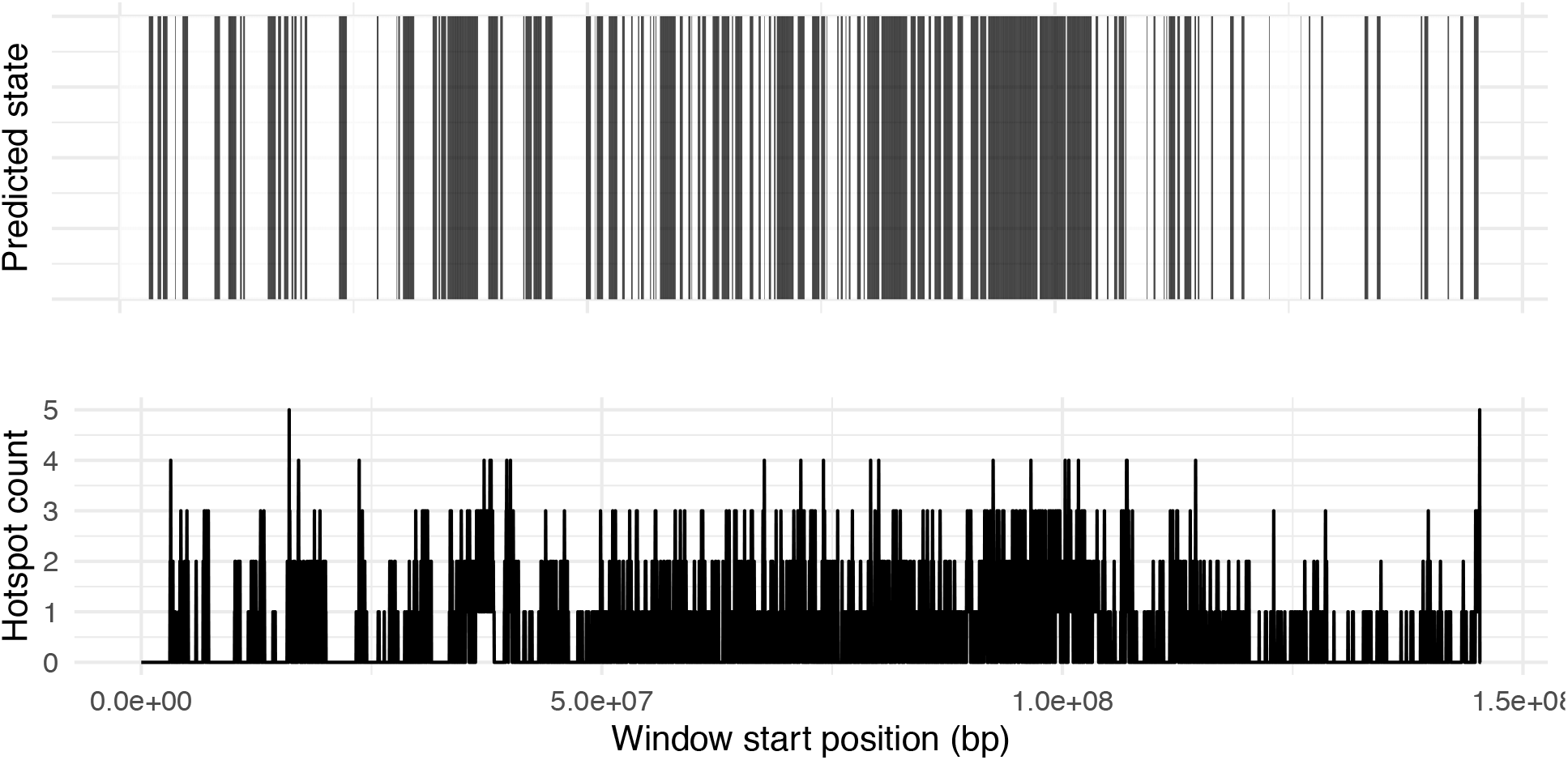
Predicted states according to best fit ZIPHMM and hotspot count for all chromosome 7 windows.

**Supplementary figure 8.**
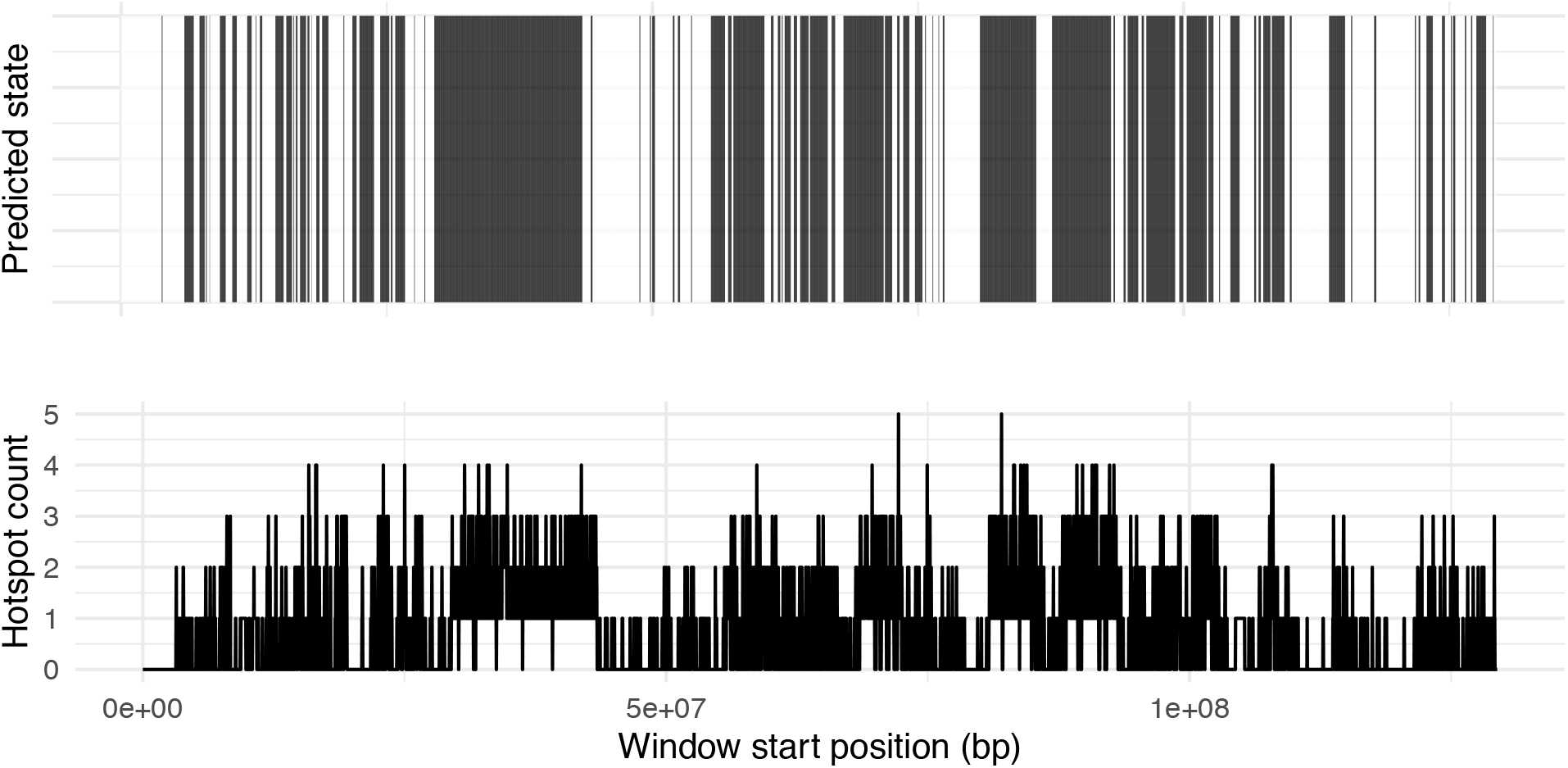
Predicted states according to best fit ZIPHMM and hotspot count for all chromosome 8 windows.

**Supplementary figure 9.**
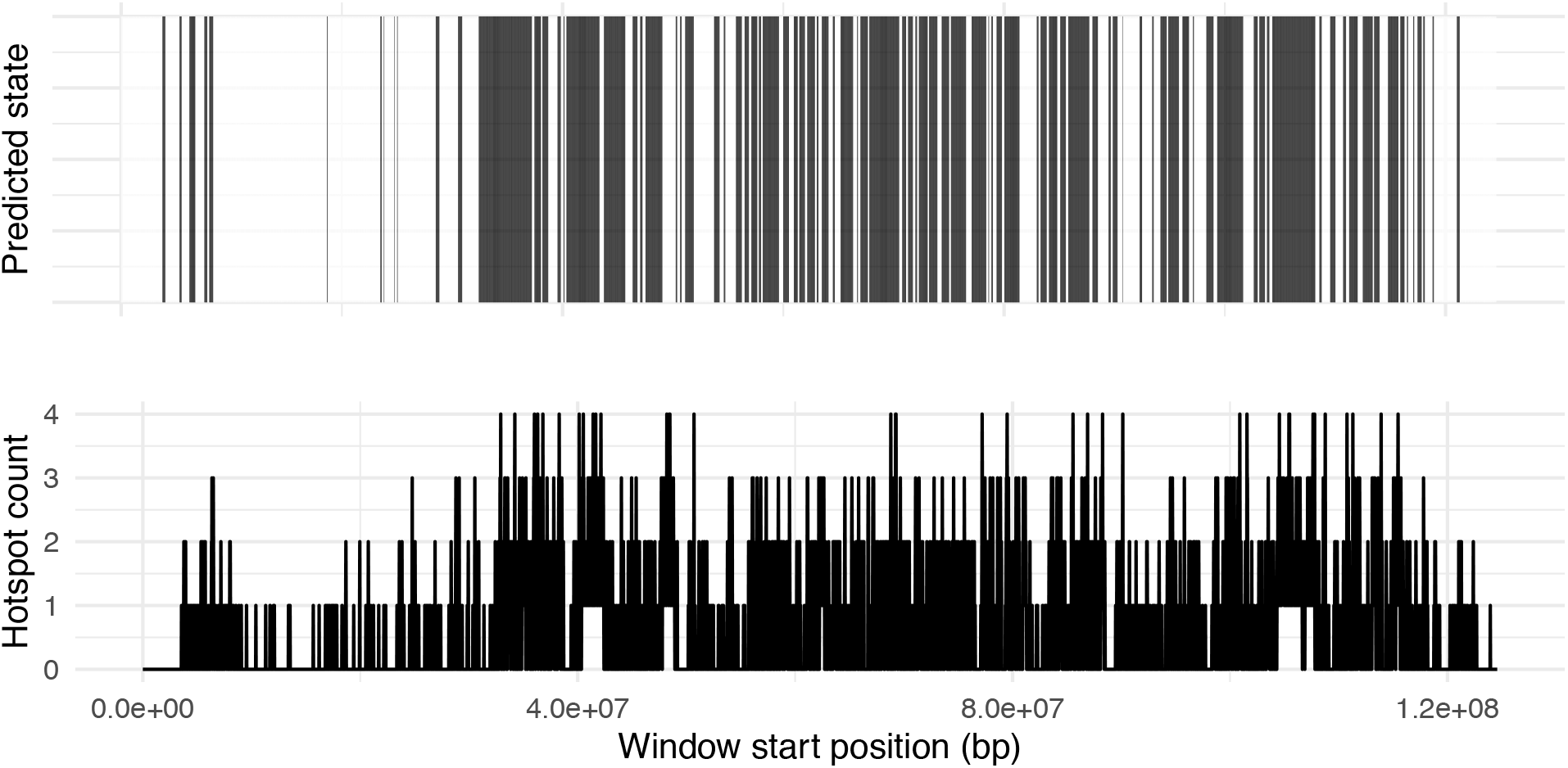
Predicted states according to best fit ZIPHMM and hotspot count for all chromosome 9 windows.

**Supplementary figure 10.**
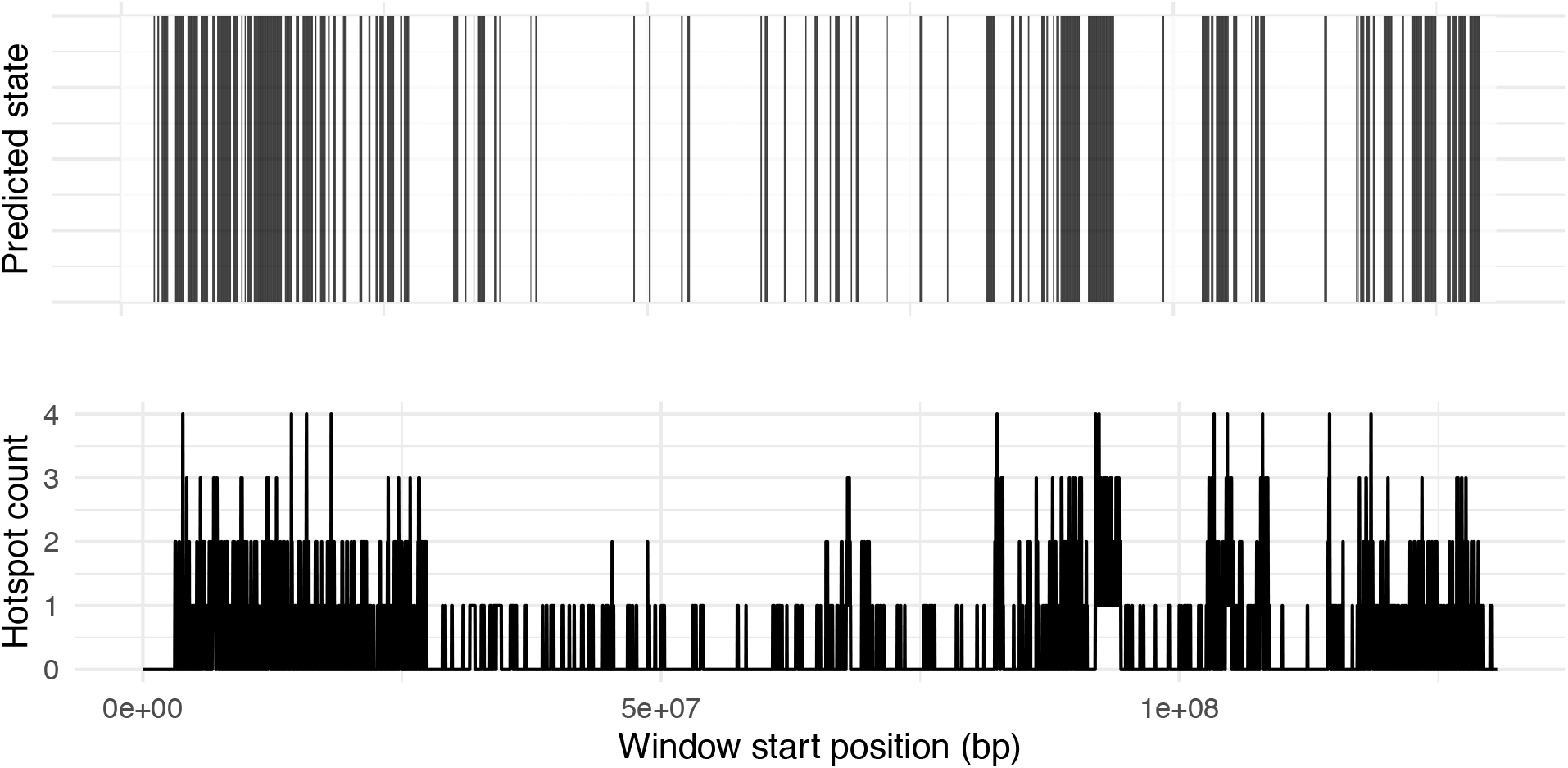
Predicted states according to best fit ZIPHMM and hotspot count for all chromosome 10 windows.

**Supplementary figure 11.**
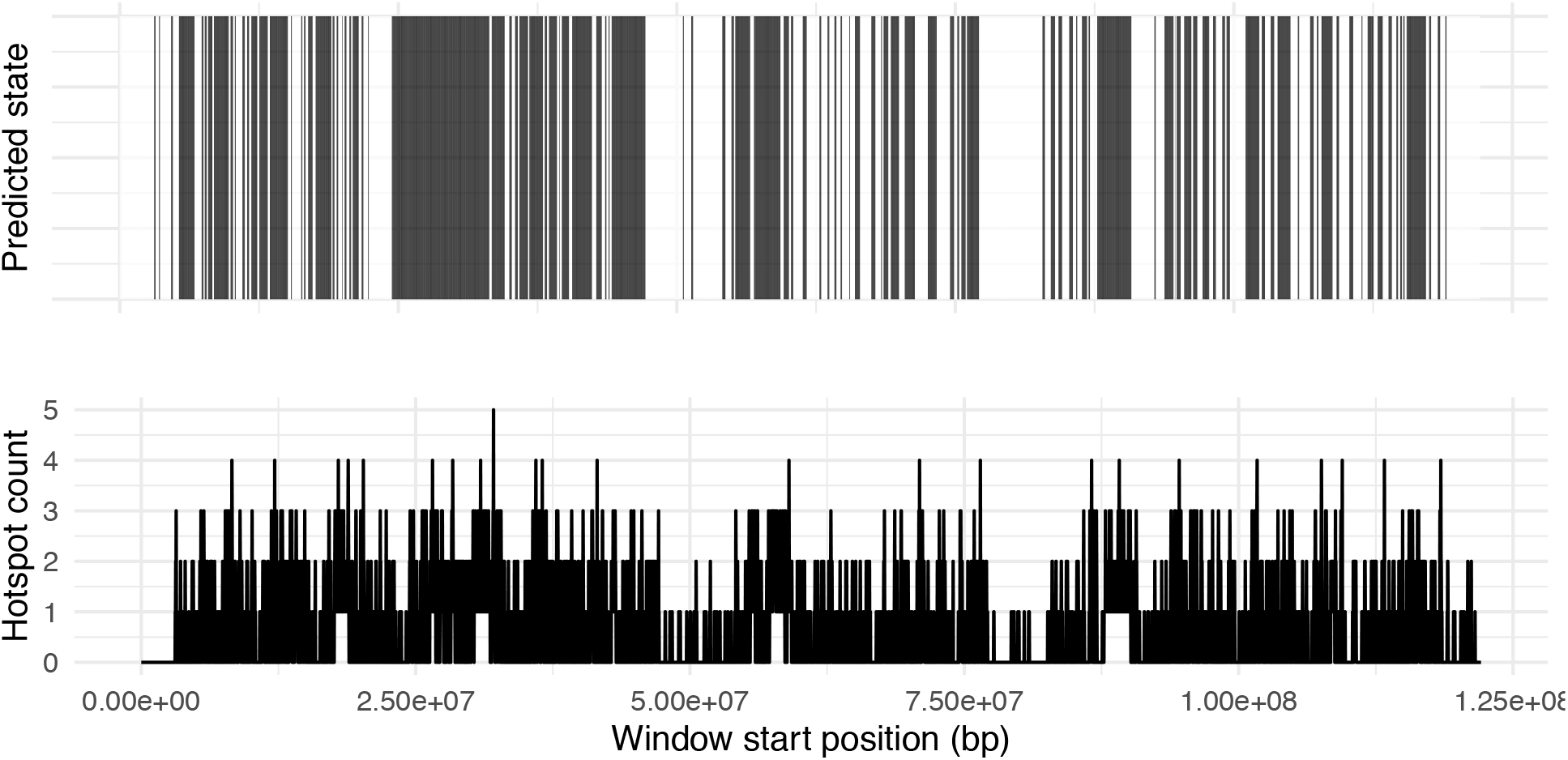
Predicted states according to best fit ZIPHMM and hotspot count for all chromosome 11 windows.

**Supplementary figure 12.**
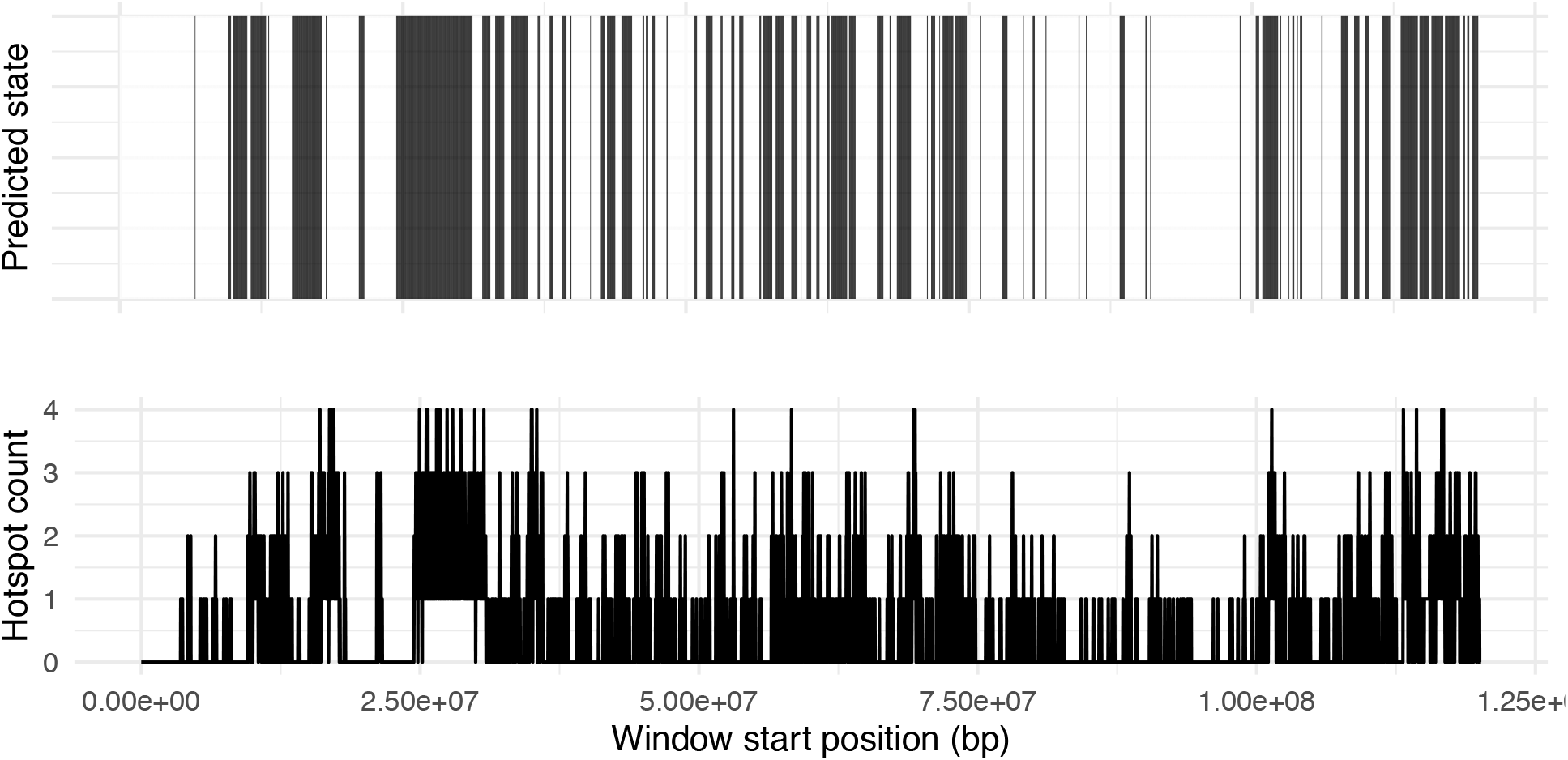
Predicted states according to best fit ZIPHMM and hotspot count for all chromosome 12 windows.

**Supplementary figure 13.**
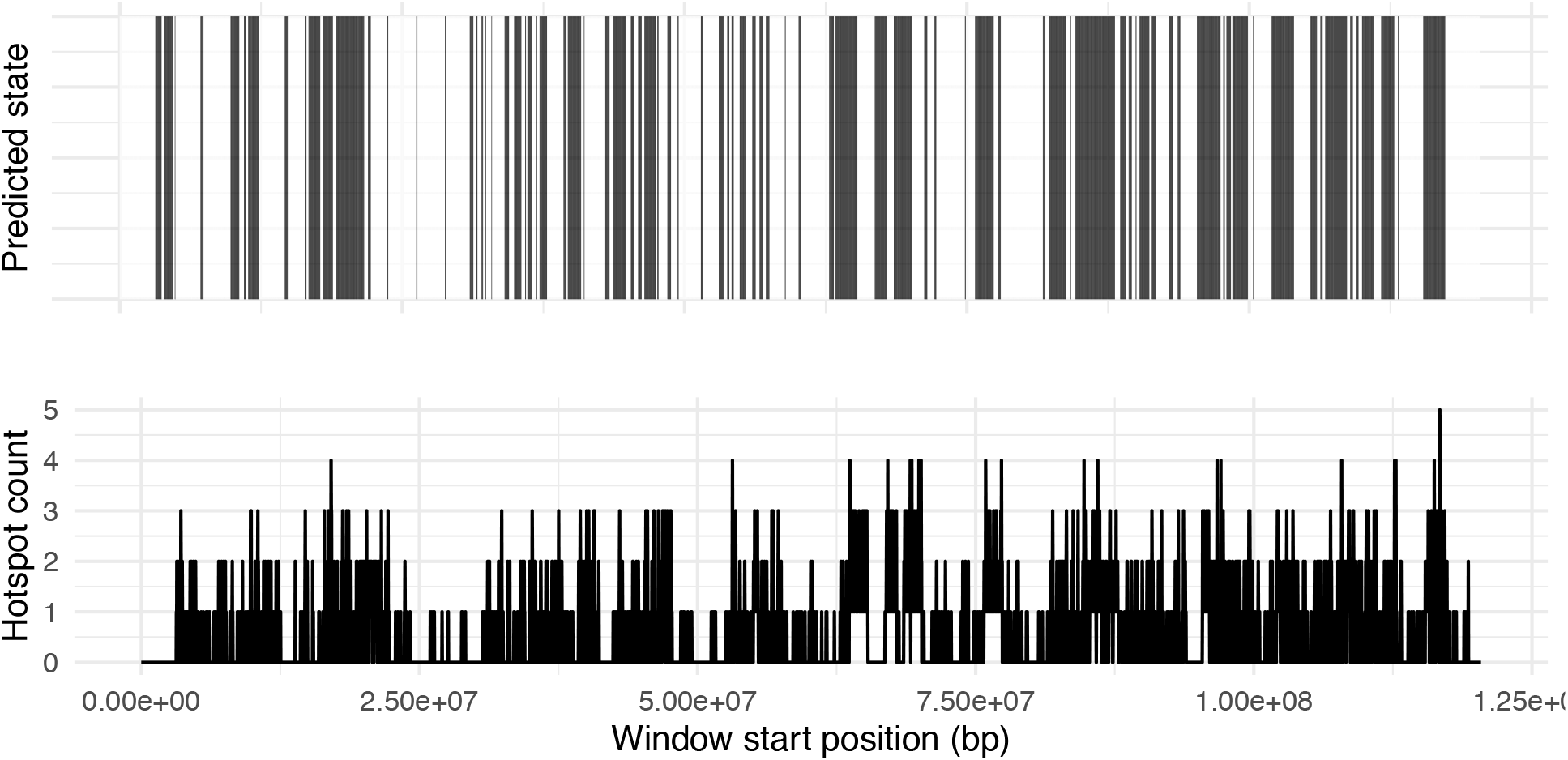
Predicted states according to best fit ZIPHMM and hotspot count for all chromosome 13 windows.

**Supplementary figure 14.**
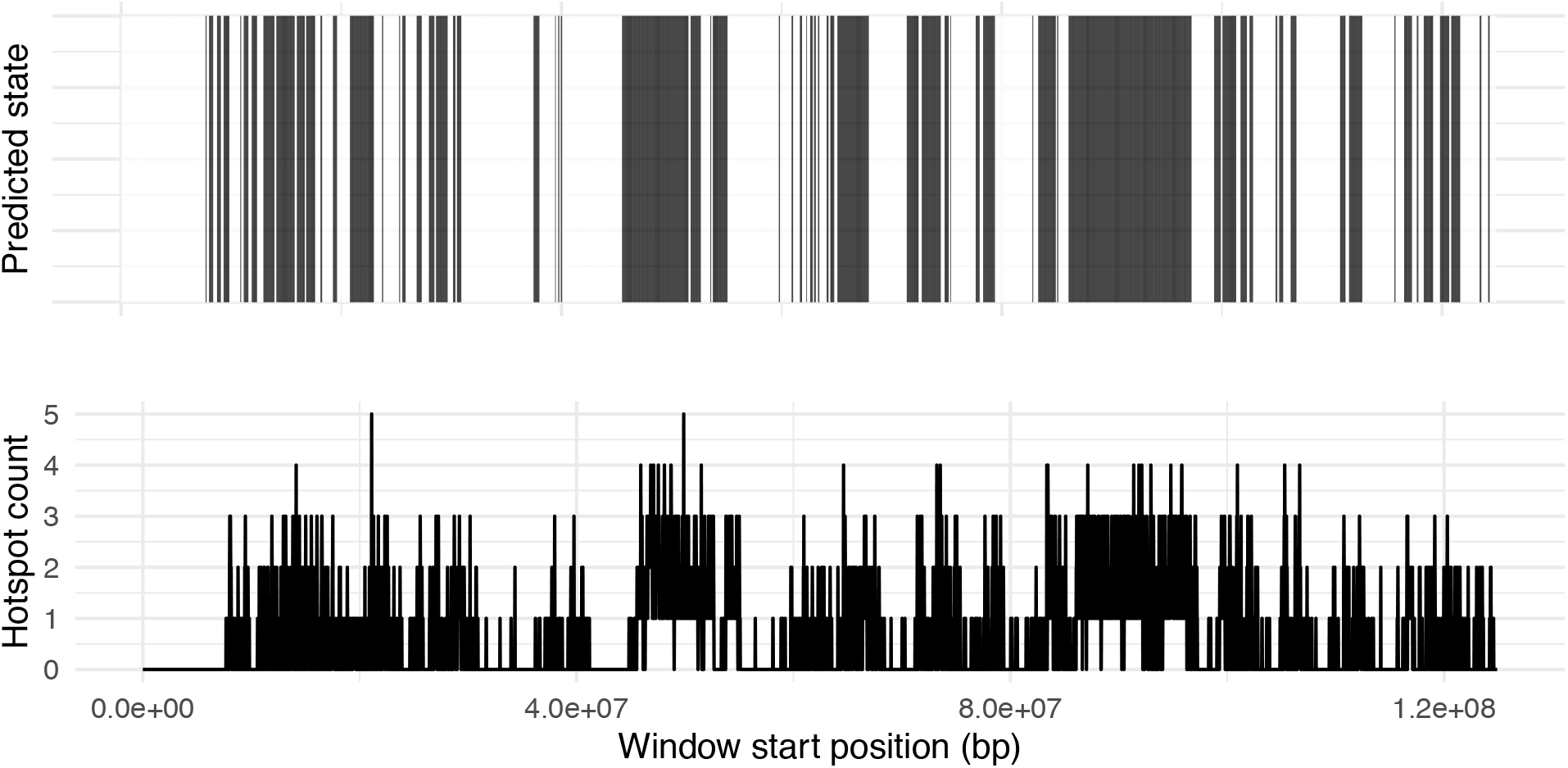
Predicted states according to best fit ZIPHMM and hotspot count for all chromosome 14 windows.

**Supplementary figure 15.**
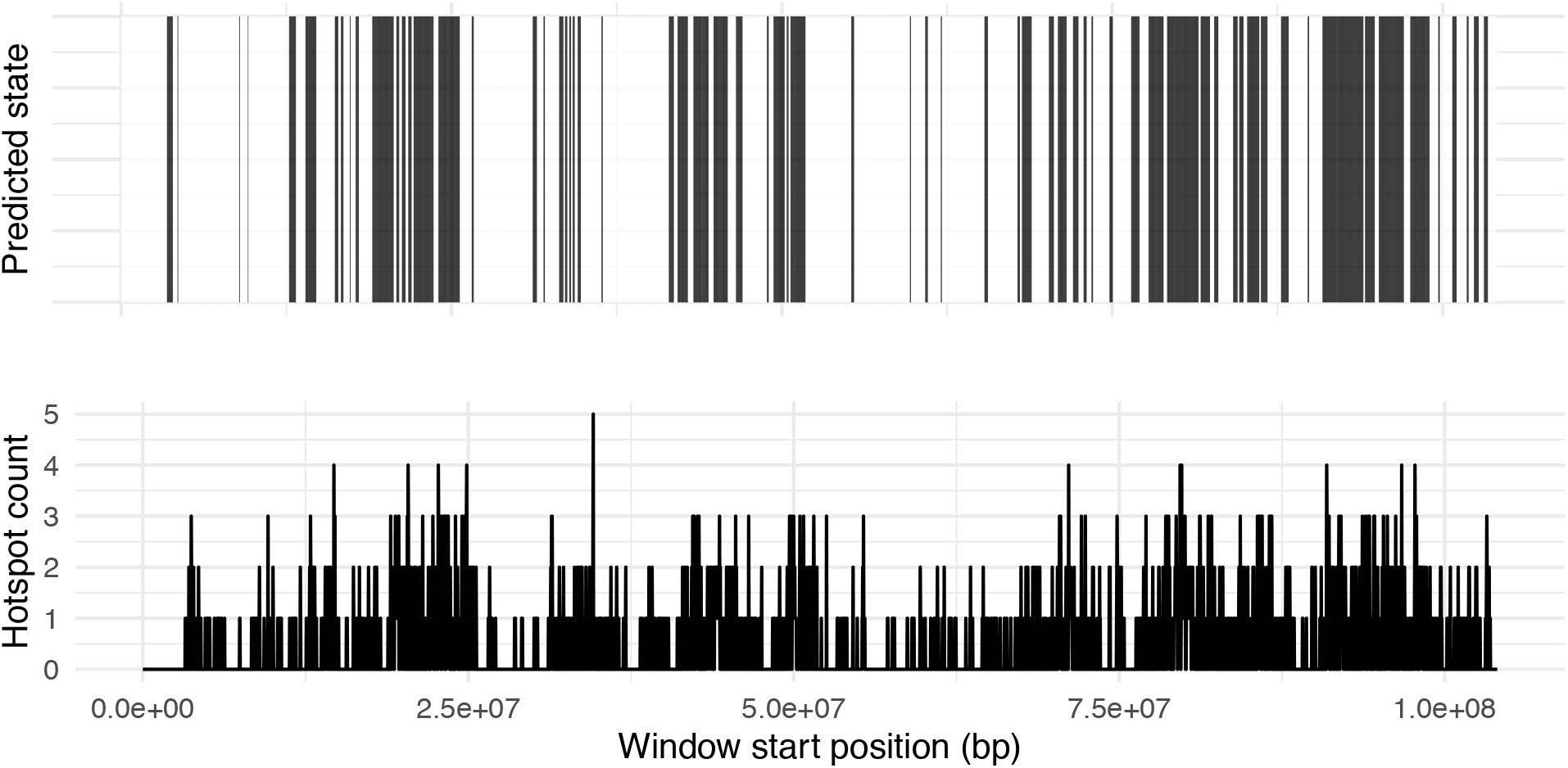
Predicted states according to best fit ZIPHMM and hotspot count for all chromosome 15 windows.

**Supplementary figure 16.**
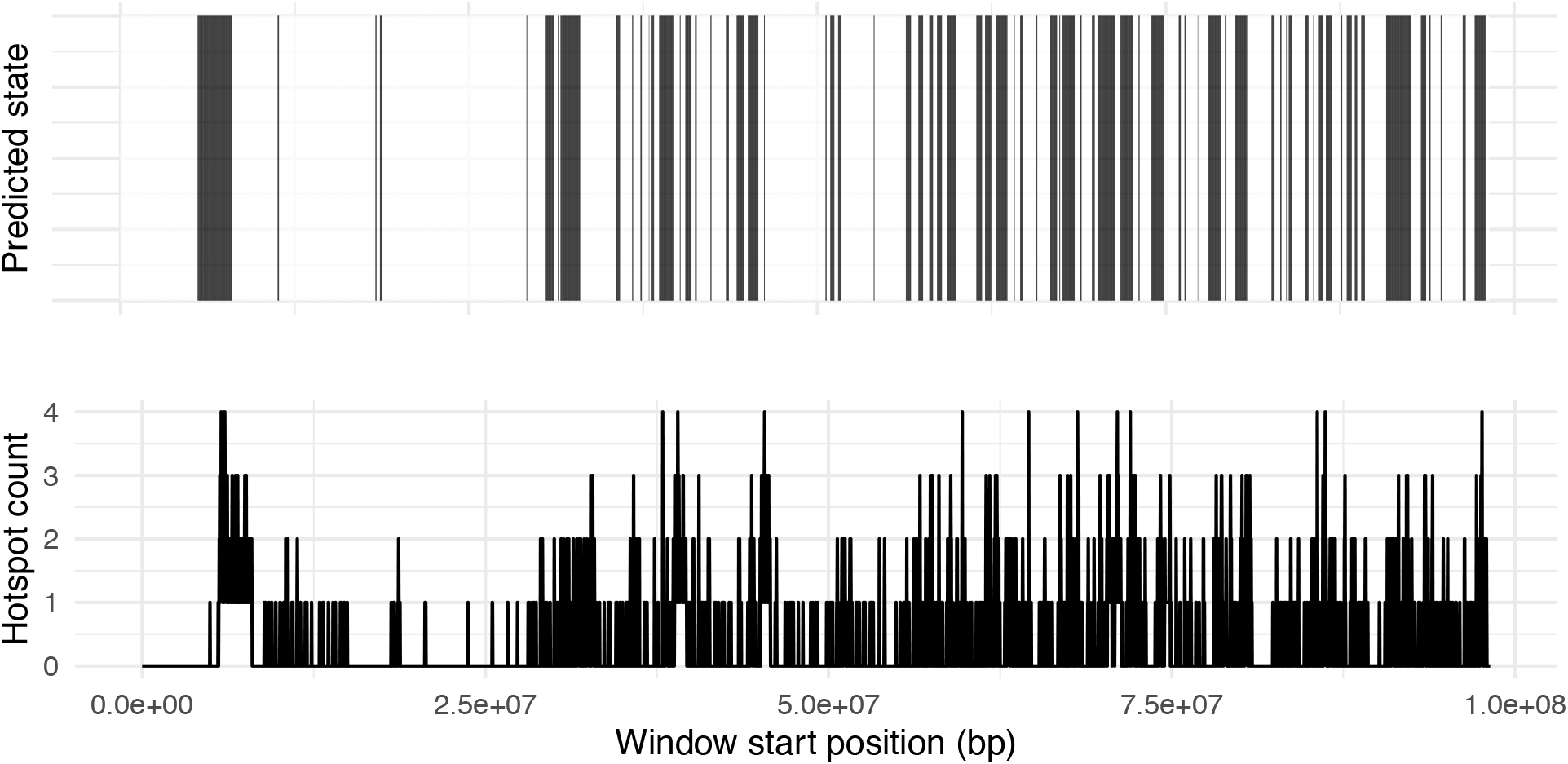
Predicted states according to best fit ZIPHMM and hotspot count for all chromosome 16 windows.

**Supplementary figure 17.**
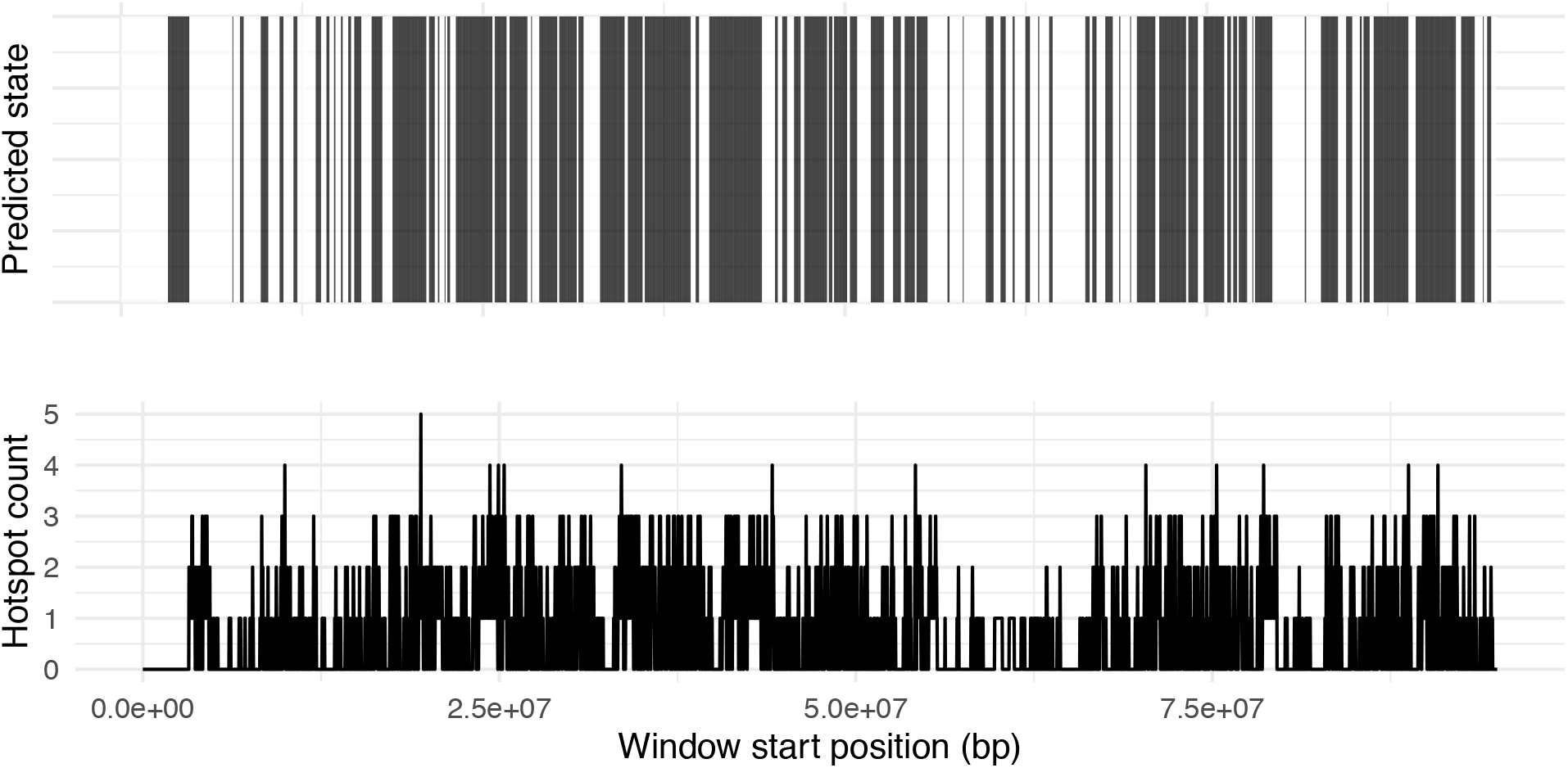
Predicted states according to best fit ZIPHMM and hotspot count for all chromosome 17 windows.

**Supplementary figure 18.**
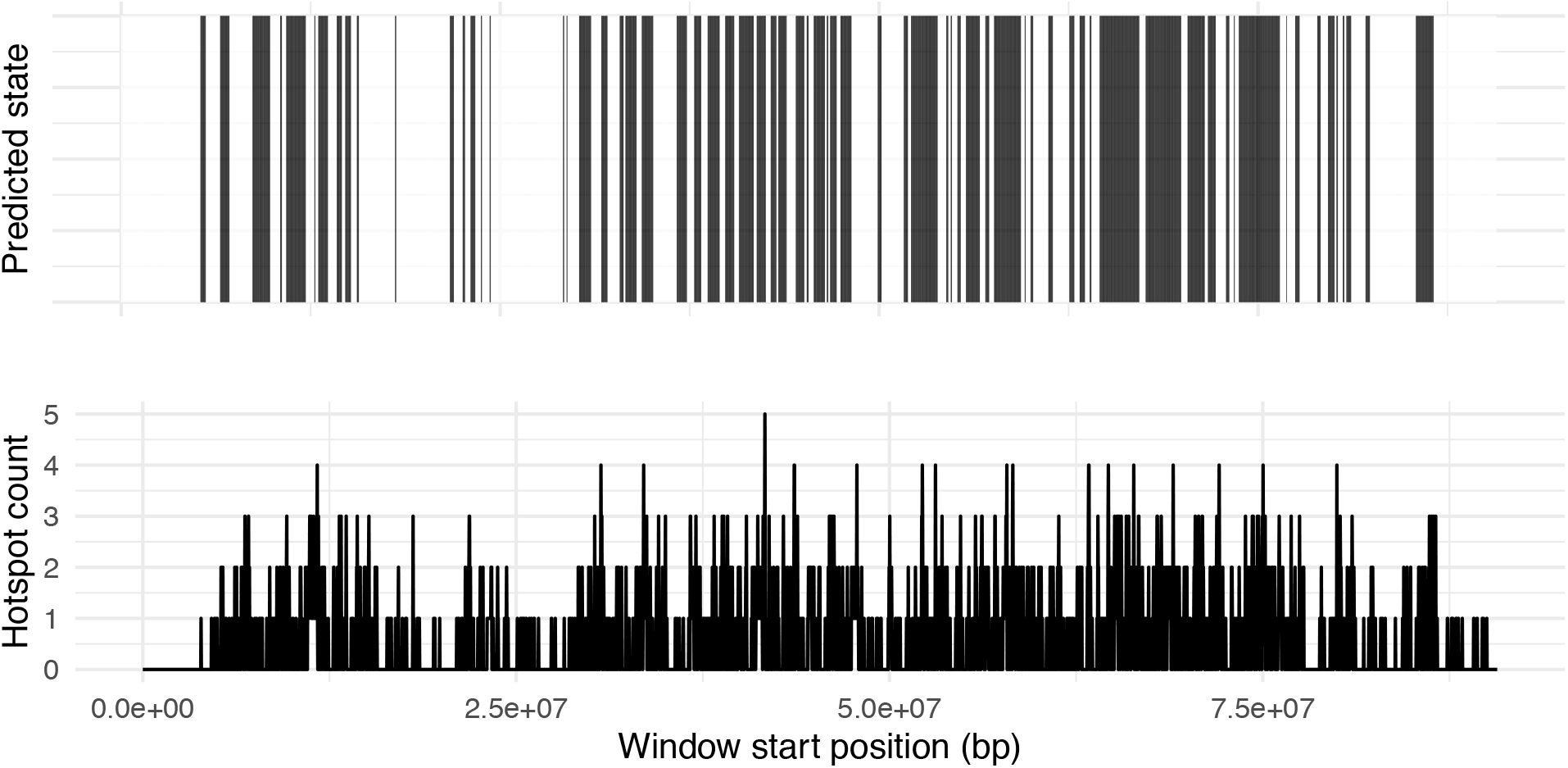
Predicted states according to best fit ZIPHMM and hotspot count for all chromosome 18 windows.

**Supplementary figure 19.**
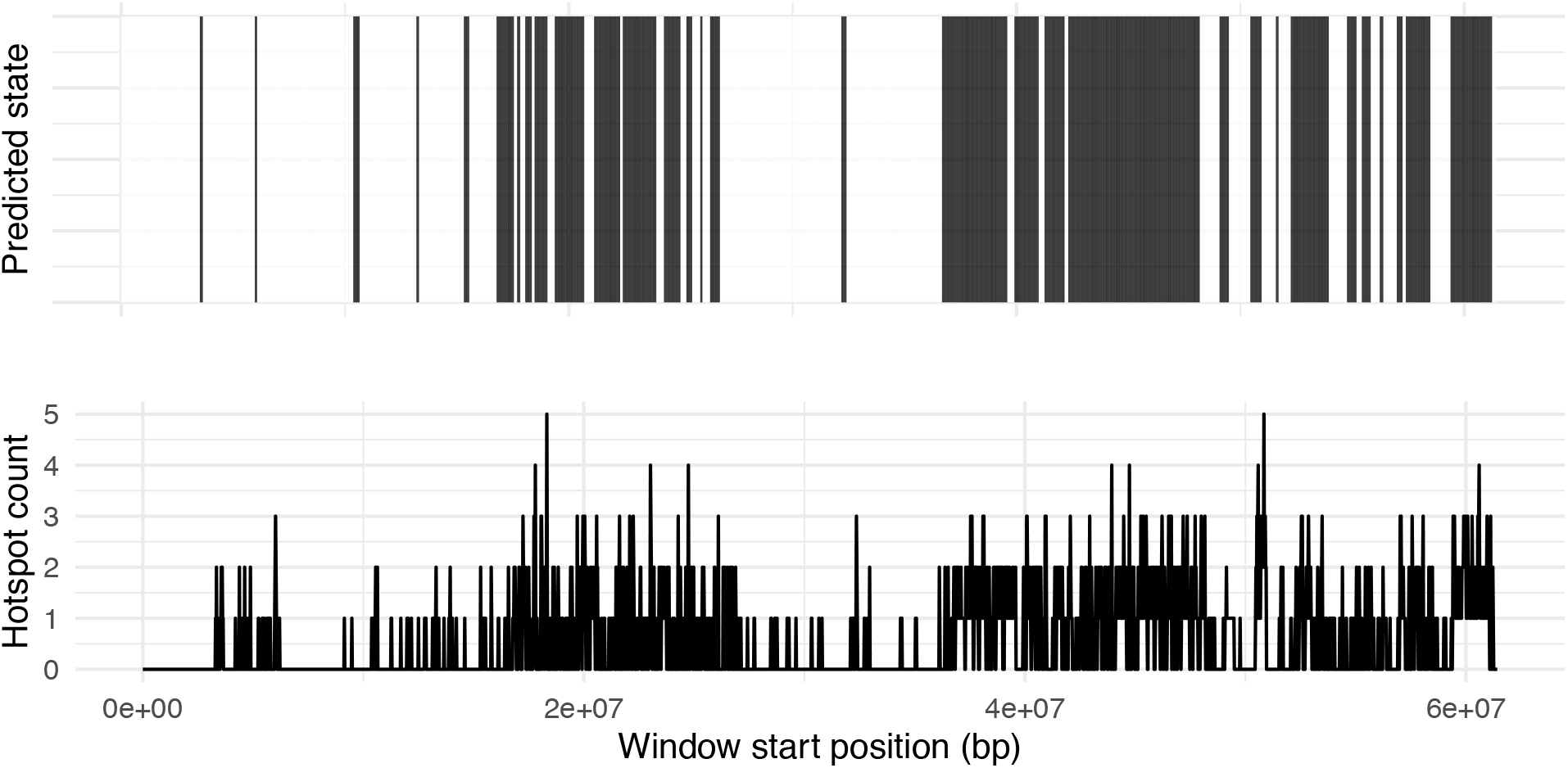
Predicted states according to best fit ZIPHMM and hotspot count for all chromosome 19 windows.

## References

Alves, I., Houle, A. A., Hussin, J. G., & Awadalla, P. (2017). The impact of recombination on human mutation load and disease. Philosophical Transactions of the Royal Society of London. Series B, Biological Sciences, 372(1736), 20160465–20160465. doi:10.1098/rstb.2016.0465

Auton, A., Fledel-Alon, A., Pfeifer, S., Venn, O., Ségurel, L., Street, T.,… Mcvean, G. (2012). A fine-scale chimpanzee genetic map from population sequencing. Science (New York, N.Y.), 336(6078), 193–198. doi:10.1126/science.1216872

Auton, A., Rui L., Ying, Kidd, J., Oliveira, K., Nadel, J., Holloway, J. K., Hayward, J. J., Cohen, P. E., Greally, J. M., Wang, J., Bustamante, C. D., & Boyko, A. R. (2013). Genetic recombination is targeted towards gene promoter regions in dogs. PLoS Genetics, 9(12), e1003984–e1003984. doi:10.1371/journal.pgen.1003984

Bai, S., Fu, K., Yin, H., Cui, Y., Yue, Q., Li, Q., Cheng, L., Tan, H., Liu, X., Guo, Y., Zhang, Y., Xie, J., He, W., Wang, Y., Feng, H., Xin, C., Zhang, J., Lin, M., Shen, B.,… Ye, L. (2018). Sox30 initiates transcription of haploid genes during late meiosis and spermiogenesis in mouse testes. Development (Cambridge), 145(13), dev164855. doi:10.1242/dev.164855

Baker, C. L., Walker, M., Kajita, S., Petkov, P. M., & Paigen, K. (2014). PRDM9 binding organizes hotspot nucleosomes and limits Holliday junction migration. Genome research, 24(5), 724–732.

Baker, Z., Schumer, M., Haba, Y., Bashkirova, L., Holland, C., Rosenthal, G. G., & Przeworski, M. (2017). Repeated losses of PRDM9-directed recombination despite the conservation of PRDM9 across vertebrates. eLife, 6. https://doi.org/10.7554/eLife.24133

Batzer, M. A., & Cordaux, R. (2009). The impact of retrotransposons on human genome evolution. Nature Reviews. Genetics, 10(10), 691–703. doi:10.1038/nrg2640

Brick, K., Smagulova, F., Khil, P., Camerini-Otero, R. D., & Petukhova, G. V. (2012). Genetic recombination is directed away from functional genomic elements in mice. Nature, 485(7400), 642.

Brunschwig, H., Levi, L., Ben-David, E., Williams, R. W., Yakir, B., & Shifman, S. (2012). Fine-scale maps of recombination rates and hotspots in the mouse genome. Genetics, 191(3), 757–764. doi:10.1534/genetics.112.141036

Buard, J., Barthès, P., Grey, C., & De Massy, B. (2009). Distinct histone modifications define initiation and repair of meiotic recombination in the mouse. The EMBO journal, 28(17), 2616–2624.

Bugge, M., Collins, A., Petersen, M. B., Fisher, J., Brandt, C., Michael Hertz, J.… & Morton, N. (1998). Non-disjunction of chromosome 18. Human molecular genetics, 7(4), 661–669. doi:10.1093/hmg/7.4.661

Campos-Sánchez, R., Cremona, M. A., Pini, A., Chiaromonte, F., & Makova, K. D. (2016). Integration and Fixation Preferences of Human and Mouse Endogenous Retroviruses Uncovered with Functional Data Analysis. PLoS Computational Biology, 12(6), e1004956–e1004956. doi:10.1371/journal.pcbi.1004956

Chen, J., Tresenrider, A., Chia, M., McSwiggen, D. T., Spedale, G., Jorgensen, V., Liao, H., van Werven, F. J., & Ünal, E. (2017). Kinetochore inactivation by expression of a repressive mRNA. eLife, 6. doi:10.7554/eLife.27417

Chen, Y., Zheng, Y., Gao, Y., Lin, Z., Yang, S., Wang, T., Wang, Q., Xie, N., Hua, R., Liu, M., Sha, J., Griswold, M. D., Li, J., Tang, F., & Tong, M. (2018). Single-cell RNA-seq uncovers dynamic processes and critical regulators in mouse spermatogenesis. Cell Research, 28(9), 879–896. doi:10.1038/s41422-018-0074-y

Choi, K., Zhao, X., Kelly, K. A., Venn, O., Higgins, J. D., Yelina, N. E., Hardcastle, T. J., Ziolkowski, P. A., Copenhaver, G. P., Franklin, F. C., McVean, G., & Henderson, I. R. (2013). Arabidopsis meiotic crossover hot spots overlap with H2A.Z nucleosomes at gene promoters. Nature genetics, 45(11), 1327–1336. doi:10.1038/ng.2766

da Cruz, I., Rodriguez-Casuriaga, R., Santieaque, F. F., Farias, J., Curti, G., Capoano, C. A.,… Geisinger, A. (2016). Transcriptome analysis of highly purified mouse spermatogenic cell populations: gene expression signatures switch from meiotic-to postmeiotic-related processes at pachytene stage. BMC Genomics, 17(281), 294. doi:10.1186/s12864-016-2618-1

Diagouraga, B., Clément, J. A.., Duret, L., Kadlec, J., de Massy, B., & Baudat, F. (2018). PRDM9 Methyltransferase Activity Is Essential for Meiotic DNA Double-Strand Break Formation at Its Binding Sites. Molecular Cell, 69(5), 853–865.e6. doi:10.1016/j.molcel.2018.01.033

Felsenstein J. (1974). The evolutionary advantage of recombination. Genetics, 78(2), 737–756.

Francis, S., Yelumalai, S., Jones, C., & Coward, K. (2014). Aberrant protamine content in sperm and consequential implications for infertility treatment. Human Fertility (Cambridge, England), 17(2), 80–89. doi:10.3109/14647273.2014.915347

Getun, I. V., Wu, Z. K., Khalil, A. M., & Bois, P. R. (2010). Nucleosome occupancy landscape and dynamics at mouse recombination hotspots. EMBO reports, 11(7), 555–560. doi:10.1038/embor.2010.79

Getun, I. V., Wu, Z. K., & Bois, P. R. (2012). Organization and roles of nucleosomes at mouse meiotic recombination hotspots. Nucleus (Austin, Tex.), 3(3), 244–250. doi:10.4161/nucl.20325

Gregorova, S., Gergelits, V., Chvatalova, I., Bhattacharyya, T., Valiskova, B., Fotopulosova, V., Jansa, P., Wiatrowska, D., & Forejt, J. (2018). Modulation of Prdm9-controlled meiotic chromosome asynapsis overrides hybrid sterility in mice. eLife, 7. doi:10.7554/eLife.34282

Grey, C., Barthès, P., Chauveau-Le Friec, G., Langa, F., Baudat, F., & De Massy, B. (2011). Mouse PRDM9 DNA-binding specificity determines sites of histone H3 lysine 4 trimethylation for initiation of meiotic recombination. PLoS biology, 9(10), e1001176.

Grey, C., Baudat, F., & de Massy, B. (2018). PRDM9, a driver of the genetic map. PLoS Genetics, 14(8), e1007479–e1007479. doi:10.1371/journal.pgen.1007479

Hassold, T., Merrill, M., Adkins, K., Freeman, S., & Sherman, S. (1995). Recombination and maternal age-dependent nondisjunction: molecular studies of trisomy 16. American journal of human genetics, 57(4), 867.

Hayashi, K., Yoshida, K., & Matsui, Y. (2005). A histone H3 methyltransferase controls epigenetic events required for meiotic prophase. Nature, 438(7066), 374–378. doi:10.1038/nature04112

Hellsten, U., Wright, K. M., Jenkins, J., Shu, S., Yuan, Y., Wessler, S. R., Schmutz, J., Willis, J. H., Rokhsar, D. S. (2013). Fine-scale variation in meiotic recombination in Mimulus inferred from population shotgun sequencing. Proceedings of the National Academy of Sciences - PNAS, 110(48), 19478–19482. doi:10.1073/pnas.1319032110

Heyer, W. D., Ehmsen, K. T., & Liu, J. (2010). Regulation of homologous recombination in eukaryotes. Annual review of genetics, 44, 113–139. doi:10.1146/annurev-genet-051710-150955

Hill, W., & Robertson, A. (1966). The effect of linkage on limits to artificial selection. Genetical Research, 8(3), 269–294. doi:10.1017/S0016672300010156

Hua, R., Wei, H., Liu, C., Zhang, Y., Liu, S., Guo, Y., Cui, Y., Zhang, X., Guo, X., Li, W., & Liu, M. (2019). FBXO47 regulates telomere-inner nuclear envelope integration by stabilizing TRF2 during meiosis. Nucleic Acids Research, 47(22), 11755–11770. doi:10.1093/nar/gkz992

Kent, W., Sugnet, C., Furey, T., Roskin, K., Pringle, T., Zahler, A., & Haussler, D. (2002). The Human Genome Browser at UCSC. Genome Research, 12(6), 996–1006. doi:10.1101/gr.229102

Kent, T. V, Uzunović, J., & Wright, S. I. (2017). Coevolution between transposable elements and recombination. Philosophical Transactions. Biological Sciences, 372(1736), 20160458. doi:10.1098/rstb.2016.0458

Lamb, N. E., Feingold, E., Savage, A., Avramopoulos, D., Freeman, S., Gu, Y.,… & Saker, D. (1997). Characterization of susceptible chiasma configurations that increase the risk for maternal nondisjunction of chromosome 21. Human molecular genetics, 6(9), 1391–1399.

Lee, K., Haugen, H. S., Clegg, C. H., & Braun, R.E. (1995). Premature Translation of Protamine 1 mRNA Causes Precocious Nuclear Condensation and Arrests Spermatid Differentiation in Mice. Proceedings of the National Academy of Sciences - PNAS, 92(26), 12451–12455. doi:10.1073/pnas.92.26.12451

Maezawa, S., Yukawa, M., Alavattam, K. G., Barski, A., & Namekawa, S. H. (2018). Dynamic reorganization of open chromatin underlies diverse transcriptomes during spermatogenesis. Nucleic Acids Research, 46(2), 593–608. doi:10.1093/nar/gkx1052

Mcvean, G. A. T., Myers, S. R., Hunt, S., Deloukas, P., Bentley, D. R., & Donnelly, P. (2004). The fine-scale structure of recombination rate variation in the human genome. Science (New York, N.Y.), 304(5670), 581–584. doi:10.1126/science.1092500

Mihola, O., Trachtulec, Z., Vlcek, C., Schimenti, J.C., & Forejt, J. (2009). A Mouse Speciation Gene Encodes a Meiotic Histone H3 Methyltransferase. Science, 323(5912), 373–375. doi:10.1126/science.1163601

Myers, S., Bottolo, L., Freeman, C., Mcvean, G., & Donnelly, P. (2005). A fine-scale map of recombination rates and hotspots across the human genome. Science (New York, N.Y.), 310(5746), 321–324. doi:10.1126/science.1117196

Myers, S., Freeman, C., Auton, A., Donnelly, P., & McVean, G. (2008). A common sequence motif associated with recombination hot spots and genome instability in humans. Nature Genetics, 40(9), 1124–1129. doi:10.1038/ng.213

Myers, S., Bowden, R., Tumian, A., Bontrop, R. E., Freeman, C., Macfie, T. S.,… Donnelly, P. (2010). Drive against hotspot motifs in primates implicates the PRDM9 gene in meiotic recombination. Science (New York, N.Y.), 327(5967), 876–879. doi:10.1126/science.1182363

Parvanov, E. D., Petkov, P. M., & Paigen, K. (2010). Prdm9 Controls Activation of Mammalian Recombination Hotspots. Science, 327(5967), 835–835. doi:10.1126/science.1181495

Powers, N. R., Parvanov, E. D., Baker, C. L., Walker, M., Petkov, P. M., & Paigen, K. (2016). The meiotic recombination activator PRDM9 trimethylates both H3K36 and H3K4 at recombination hotspots in vivo. PLoS genetics, 12(6), e1006146.

Quinlan, A. R., & Hall, I. M. (2010). BEDTools: a flexible suite of utilities for comparing genomic features. Bioinformatics, 26(6), 841–842. doi:10.1093/bioinformatics/btq033

R Core Team (2019). R: A language and environment for statistical computing. R Foundation for Statistical Computing, Vienna, Austria. URL https://www.R-project.org/.

Robinson, W. P., Kuchinka, B. D., Bernasconi, F., Petersen, M. B., Schulze, A., Brøndum-Nielsen, K.,… & Schuffenhauer, S. (1998). Maternal meiosis I non-disjunction of chromosome 15: dependence of the maternal age effect on level of recombination. Human molecular genetics, 7(6), 1011–1019.

Schield, Drew R, Pasquesi, Giulia I M, Perry, Blair W, Adams, Richard H, Nikolakis, Zachary L, Westfall, Aundrea K, Orton, Richard W, Meik, Jesse M, Mackessy, Stephen P, & Castoe, Todd A. (2020). Snake Recombination Landscapes Are Concentrated in Functional Regions despite PRDM9. Molecular Biology and Evolution, 37(5), 1272–1294. doi:10.1093/molbev/msaa003

Schwarzkopf, E. J., Motamayor, J. C., & Cornejo, O. E. (2020). Genetic differentiation and intrinsic genomic features explain variation in recombination hotspots among cocoa tree populations. BMC Genomics, 21(1), 332–332. doi:10.1186/s12864-020-6746-2

Singhal, S., Leffler, E. M., Sannareddy, K., Turner, I., Venn, O., Hooper, D. M.,… Przeworski, M. (2015). Stable recombination hotspots in birds. Science (New York, N.Y.), 350(6263), 928–932. doi:10.1126/science.aad0843

Smagulova, F., Gregoretti, I. V., Brick, K., Khil, P., Camerini-Otero, R. D., & Petukhova, G. V. (2011). Genome-wide analysis reveals novel molecular features of mouse recombination hotspots. Nature, 472(7343), 375.

Smit A. F. (1996). The origin of interspersed repeats in the human genome. Current opinion in genetics & development, 6(6), 743–748. doi:10.1016/s0959-437x(96)80030-x

Smit, AFA, Hubley, R & Green, P. RepeatMasker Open-4.0. 2013-2015 <http://www.repeatmasker.org>.

Stevison, L. S., Woerner, A. E., Kidd, J. M., Kelley, J. L., Veeramah, K. R., McManus, K. F.,… Wall, J. D. (2016). The Time Scale of Recombination Rate Evolution in Great Apes. Molecular Biology and Evolution, 33(4), 928–945. doi:10.1093/molbev/msv331

Thomas, N. S., Ennis, S., Sharp, A. J., Durkie, M., Hassold, T. J., Collins, A. R., & Jacobs, P. A. (2001). Maternal sex chromosome non-disjunction: evidence for X chromosome-specific risk factors. Human molecular genetics, 10(3), 243–250.

Tseden, K., Topaloglu, Ö., Meinhardt, A., Dev, A., Adham, I., Müller, C., Wolf, S., Böhm, D., Schlüter, G., Engel, W., & Nayernia, K. (2007). Premature translation of transition protein 2 mRNA causes sperm abnormalities and male infertility. Molecular Reproduction and Development, 74(3), 273–279. doi:10.1002/mrd.20570

Walker, M., Billings, T., Baker, C. L., Powers, N., Tian, H., Saxl, R. L.,… Petkov, P. M. (2015). Affinity-seq detects genome-wide PRDM9 binding sites and reveals the impact of prior chromatin modifications on mammalian recombination hotspot usage. Epigenetics & Chromatin, 8(1), 31–31. doi:10.1186/s13072-015-0024-6

Xu, Z. and Liu, Y. (2018). ziphsmm: Zero-Inflated Poisson Hidden (Semi-)Markov Models. R package version 2.0.6. https://CRAN.R-project.org/package=ziphsmm

Zeileis, A., Kleiber, C., & Jackman, S. (2008). Regression Models for Count Data in R. Journal of Statistical Software, 27(8). doi:10.18637/jss.v027.i08

